# Phenotypic and functional characterisation of first trimester human placental macrophages, Hofbauer cells

**DOI:** 10.1101/2020.09.03.279919

**Authors:** Jake Thomas, Anna Appios, Xiaohui Zhao, Roksana Dutkiewicz, Maria Donde, Colin Lee, Praveena Naidu, Christopher Lee, Joana Cerveira, Bing Liu, Florent Ginhoux, Graham Burton, Russell S. Hamilton, Ashley Moffett, Andrew Sharkey, Naomi McGovern

## Abstract

Hofbauer cells (HBC) are a population of macrophages found in high abundance within the stroma of the first trimester human placenta. HBC are the only fetal immune cell population within the stroma of healthy placenta. However, the functional properties of these cells are poorly described. Aligning with their predicted origin via primitive haematopoiesis, we find that HBC are transcriptionally similar to yolk sac macrophages. Phenotypically, HBC can be identified as HLA-DR^-^FOLR2^+^ macrophages. We identify a number of factors HBC secrete (including IL-8 and MMP-9) that could affect placental angiogenesis and remodelling. We determine that HBC have the capacity to play a defensive role, where they are responsive to Toll-like receptor stimulation and are microbicidal. Finally, we also identify a population of placenta-associated maternal macrophages (PAMM1a) that adhere to the placental surface and express factors, such as fibronectin, that may aid in repair.

**Summary:** Using transcriptomic and proteomic data, Thomas, J. et al, analyse human first trimester placental macrophages and delineate markers that identify them. They also reveal that Hofbauer cells have microbicidal capacity, providing the fetus with an additional layer of protection from certain microbes.

**One sentence summary:** Hofbauer cells are primitive placental macrophages with a unique phenotype and role in fetal defence.

**Non-standard abbreviation:** Placental associated maternal monocytes/macrophages, PAMM.

## Introduction

Macrophages are found within all human tissues where, within the adult, they mediate tissue homeostasis, development, repair and immunity. During embryonic development the first macrophages to seed all tissues are derived through a process called primitive haematopoiesis. These macrophages, commonly termed ‘primitive’ macrophages, are distinct to those generated through definitive haematopoiesis as there is no monocyte intermediate(Ginhoux et al., 2010; Gomez Perdiguero et al., 2015). Although in some species such as the mouse, primitive haematopoiesis is thought to only occur within the yolk sac (YS), during human embryonic development primitive haematopoiesis also takes place in the placenta(Van Handel et al., 2010).

The placenta is a major organ that regulates the health of both the mother and developing fetus during pregnancy. The human placenta develops from the trophoectoderm, the outer layer of the pre-implantation blastocyst, which forms at ~5 days post fertilisation (dpf)(Turco and Moffett, 2019). As the placenta develops, highly branched villous tree-like structures form, which contain fibroblasts, immature capillaries and macrophages, termed Hofbauer cells (HBC) (**Figure 1A**). The mesenchymal core is surrounded by a bilayer of specialized placental epithelial cells called trophoblast. The outermost syncytiotrophoblast (SCT) layer, in contact with maternal blood, is formed by fusion of underlying cytotrophoblast cells(Turco and Moffett, 2019). HBC have been identified within the placenta around day 18 post-conception(Castellucci et al., 1987; Boyd et al., 1970), before the placenta is connected to the embryonic circulation(Van Handel et al., 2010).

**Figure 1.**
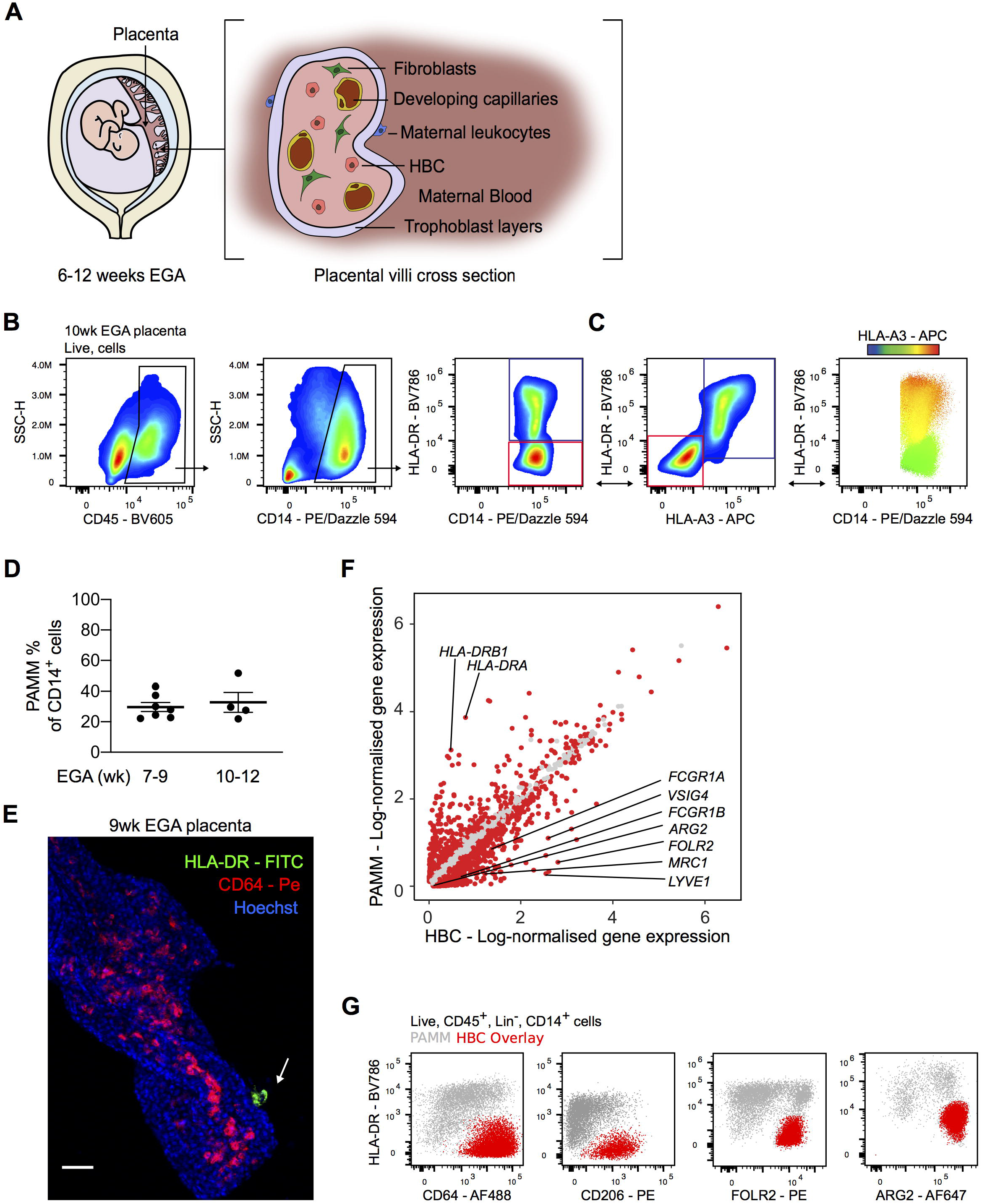
Anti-HLA antibodies allow for the specific identification of Hofbauer cells by flow cytometry. (A) Schematic drawing of the human placenta and a villous cross-section. HBC – Hofbauer cells, EGA – Estimated gestational age. (B) Representative flow cytometric gating strategy identifying two placental macrophage populations based on HLA-DR expression. Blue gate – HLA-DR^+^ macrophages. Red gate – HLA-DR^-^ macrophages. (C) Differential expression of HLA-A3 within the CD14^+^ macrophage gate, shown by biaxial plot and heatmap overlay. Maternal macrophages are indicated by the blue gate (HLA-DR^+^HLA-A3^+^), fetal macrophages are indicated by the red gate (HLA-DR^-^HLA-A3^-^). Bidirectional arrows depict equivalent cells. (D) Quantification of the abundance of PAMM within CD14^+^ placental cell suspensions across the indicated EGA. Each data point indicates a separate donor (n=11). (E) Whole-mount immunofluorescence of a placental villus, where HBC stained with CD64 (red) are within villous stroma, and PAMM stained with HLA-DR (green, white arrow) are on the syncytial layer. Cell nuclei are stained with Hoechst (blue). Scale bar = 50μm. Representative image of n=3. (F) Scatterplot showing log-normalised gene expression of HBC (x-axis) and PAMM (y-axis) clusters derived from scRNAseq data analysis. Red dots represent genes that are differentially expressed with an adjusted p value <0.01 (Wilcox rank sum test). (G) Flow cytometric analysis of expression of indicated markers by HBC (identified with anti-HLA antibodies in red overlay) and PAMM (grey). Representative plots of n=3. Data are represented as mean ±SEM (D).

A number of recent studies have profiled the gene expression of human embryonic macrophage populations(Stewart et al., 2019; Vento-Tormo et al., 2018). However, studies demonstrating their functional properties remain limited. Our previous work demonstrating that 2^nd^ trimester fetal dendritic cells are functionally active and responsive to TLR stimulation(McGovern et al., 2017) led us to query if primitive macrophages have similar capabilities. In particular, we were interested in determining if HBC demonstrate microbicidal capacity, as they are the only fetal immune cells found within the stroma of the human placenta, the crucial tissue barrier-site between maternal tissues and the fetus.

In this study we sought to develop a technique that would allow us to characterise the properties of HBC isolated from first trimester human placentas. Using a novel flow cytometric gating strategy, we find that commonly used protocols for the isolation of HBC from first trimester placentas yield a heterogenous population of macrophages that also consist of placenta-associated maternal monocyte/macrophage (PAMM) subsets. We demonstrate that HBC have a unique phenotype, specific to the placental niche; they do not express HLA-DR and highly express folate receptor 2 (FOLR2). We identify a range of factors HBC secrete that possibly affect placental angiogenesis and remodelling, including IL-8, osteopontin and MMP-9. We show that HBC are responsive to TLR stimulation and do have microbicidal capacity, and can thereby play a defensive role for the fetus. Finally, we identify a novel population of placenta-associated maternal macrophages (PAMM1a), that could function in tissue repair. Our findings provide novel insights into the properties of human primitive macrophages, and the roles of HBC in placental homeostasis.

## Results

### Identification of Hofbauer cells using anti-HLA antibodies

Previous reports phenotyping HBC isolated from the placenta have yielded conflicting results(Sutton et al., 1983; Böckle et al., 2008; Bulmer et al., 1988; Goldstein et al., 1988; Reyes and Golos, 2018). We first sought to determine the true identity of first trimester HBC using multi-parameter flow cytometry. Employing a commonly used protocol to isolate HBC(Tang et al., 2011) (**Figure S1A**), we obtain a CD45^+^CD14^+^ macrophage population that is heterogeneous for HLA-DR expression (**Figure 1B**). We sought to determine if the observed heterogeneity is due to maternal monocyte/macrophage populations contaminating the HBC placental isolates.

Fetal cells express both maternally and paternally derived genes. To determine if maternal cells contaminate the HBC preparations and contribute to the heterogeneity in observed HLA-DR expression, we added antibodies to common HLA allotypes (HLA-A3, HLA-B7 and HLA-A2) to our flow cytometry panel. The specificity of these antibodies was previously verified by quantitative polymerase chain reaction (qPCR) on DNA from blood samples of HLA-typed donors. We chose to use anti-HLA antibodies instead of sex chromatin staining (to identify male fetal cells), so that we could develop a flow cytometry panel to allow isolation of live cells for functional assays. By using anti-HLA antibodies and analysing matched maternal blood or decidual cells (**Figure S1D, S1E)**, we consistently observe that the variable population of HLA-DR^+^ cells in first trimester placental digests are maternal in origin (**Figure 1C, S1A-C**). We termed the maternal cells obtained in placenta digests ‘placenta-associated maternal monocytes/macrophages’ (PAMM).

We had expected that maternal contamination of placental macrophage cell isolates would only become significant from the 10^th^ week of gestation, the time when maternal blood flow to the intervillous space is fully established(Burton et al., 2009b). However, application of our new gating strategy to placental digests of 7-9 wk estimated gestational age (EGA) and 10-12 wk EGA, demonstrated that maternal cells make a significant contribution to the CD14^+^ macrophage populations isolated from the placenta as early as the 7^th^ week of gestation, comprising 20-40% of CD14^+^ cells (**Figure 1D**). Whole-mount immunofluorescence microscopy revealed that while HLA-DR^+^ cells do adhere to the SCT layer of the placental villi, HLA-DR^+^ cells are not present within the stromal core from the 7^th^ to 10^th^ week of gestation (**Figure 1E)**. These findings are in line with recent single cell RNA sequencing (scRNAseq) studies of placental cell isolates(Tsang et al., 2017; Vento-Tormo et al., 2018) where maternal cells were also observed. Our data demonstrate that maternal cell contamination is higher than has previously been appreciated within first trimester placental cell suspensions. These findings will have a significant impact on *in vitro* studies that aim to determine the specific functional properties of HBC.

### Identification of specific markers for HBC

A limitation of using anti-HLA antibodies is that they can only distinguish maternal and fetal cells where there is a maternal/fetal HLA-mismatch for these specific allotypes. We therefore sought to identify markers that would allow us to confidently distinguish maternal from fetal cells independently of HLA antibodies. To do this, we carried out transcriptomic analysis of first trimester placental cells using a publicly available scRNAseq dataset(Vento-Tormo et al., 2018). Clustering and uniform manifold approximation and projection (UMAP) visualisation of 22,618 placental single cells identifies 2 distinct macrophage populations, as indicated by *CD68* expression (**Figure S1F**, **S1G)**. Consistent with our flow cytometry analysis, PAMM can be readily identified as HLA-DR^hi^, while HBC are HLA-DR^-^ (**Figure SIG**). Expression of male *(RPS4Y1)* and female *(XIST)* specific genes in placental cells from male fetal donors confirms the fetal and maternal origin of HBC and PAMM clusters (**Figure S1H**).

A total of 962 genes are significantly differentially expressed genes (DEGs) (adjusted p value < 0.01) between PAMM and HBC clusters. *FCGR1A, FCGR1B, VSIG4, MRC1, FOLR2* and *LYVE-1* are upregulated within HBC. Unlike adult macrophages, HBC do not express HLA-DR. In contrast, PAMM highly express *HLA-DRB1* and *HLA-DRA* (**Figure 1F**). We tested a number of additional HBC specific markers that were identified by analysis of the sequencing data, including FOLR2, CD64 and CD206. We also analysed the expression of arginase 2, which has previously been shown to be expressed by fetal immune cells(McGovern et al., 2017). Samples from donors where the anti-HLA antibodies distinguished maternal from fetal cells were used. These markers are expressed by HBC at the protein level by flow cytometry (**Figure 1G**). Any of these 4 markers, in combination with HLA-DR, allows us to confidently distinguish HBC from PAMM within first trimester samples. The combination of FOLR2 and HLA-DR provides the clearest separation for the isolation of HBC.

### Hofbauer cells are transcriptionally similar to ‘primitive’ macrophages and proliferate *in situ*

HBC are predicted to be ‘primitive’ macrophages derived directly from progenitors independent of monocytes. A recent study characterising the transcriptional landscape of human macrophage development, identified a population of true primitive yolk sac macrophages (YS_Mac1) from a Carnegie stage 11 embryo (~4 weeks post conception)(Bian et al., 2020). Consistent with their predicted primitive origins, HBC are enriched for a gene signature derived from YS_Mac1, but not embryonic monocytes (**Figure 2A, Supplementary File 1**). Integration of first trimester placental(Vento-Tormo et al., 2018) and early human fetal myeloid scRNAseq(Bian et al., 2020) datasets reveals a high degree of transcriptional similarity between HBC and primitive YS_Mac1 (**Figure 2B, 2C)**. PAMM however, display transcriptional similarity to embryonic monocytes, reflecting their monocytic origins (**Figure 2A, 2C)**.

**Figure 2.**
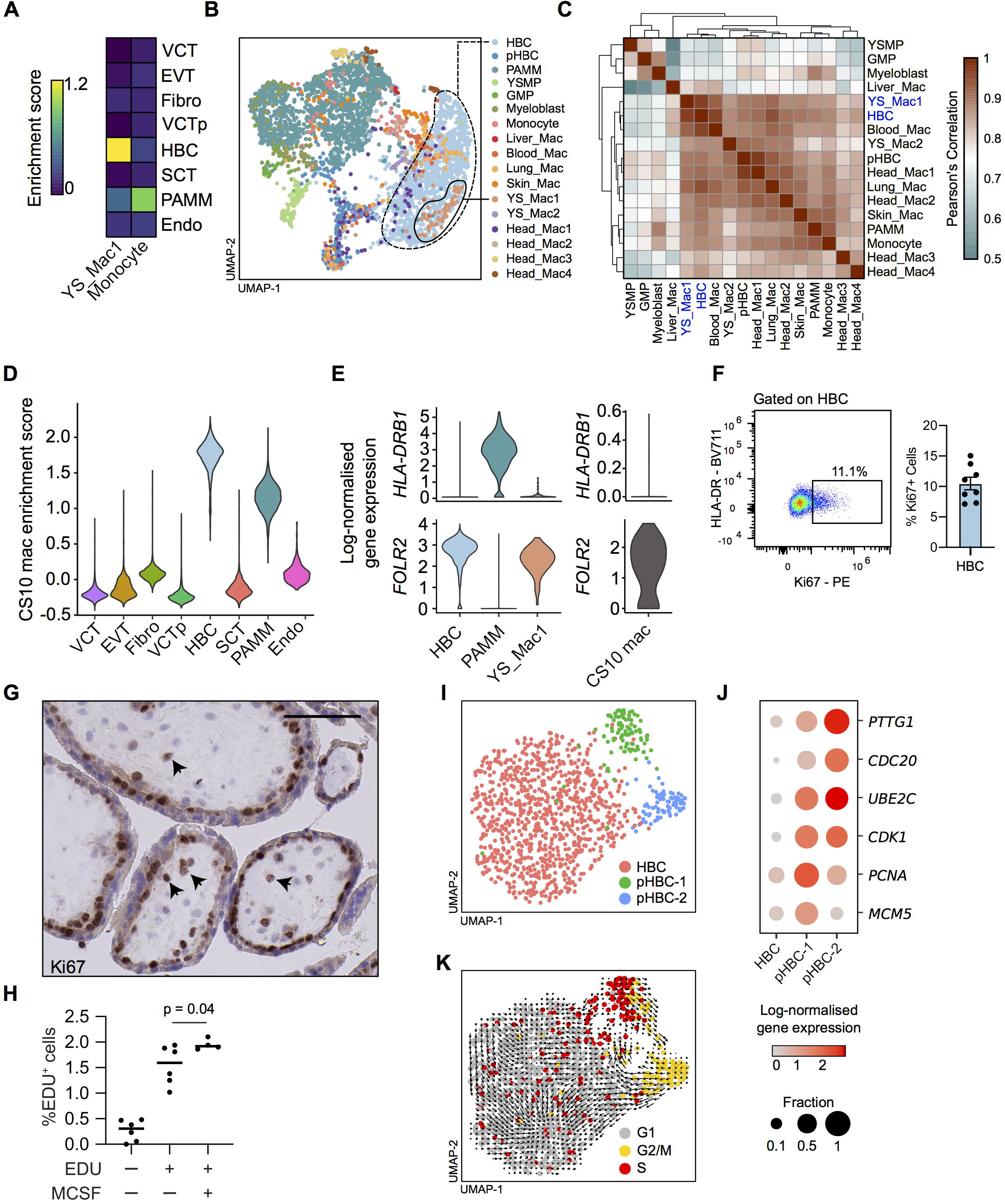
First trimester HBC are transcriptionally similar to ‘primitive’ macrophages and proliferate *in situ.* (A) Heatmap of placental scRNAseq cluster mean enrichment scores for extra-embryonic yolk sac (YS) macrophage and embryonic monocyte gene signatures(Bian et al., 2020). HBC – Hofbauer cell, PAMM – placenta-associated maternal monocytes/macrophages, VCT – villous cytotrophoblast, VCTp – proliferating villous cytotrophoblast, SCT – syncytiotrophoblast, EVT – Extravillous trophoblast, Fibro – Fibroblasts, Endo – Endothelial cells. (B) UMAP visualisation of 3,846 single cell transcriptomes from first trimester placenta and embryonic myeloid cells(Bian et al., 2020). pHBC – proliferating HBC, YSMP – yolk sac-derived myeloid-biased progenitors, GMP – granulocyte–monocyte progenitors, YS_Mac – yolk sac macrophage. (C) Heatmap depicting transcriptomic similarity between annotated clusters. Clusters are ordered according to hierarchical clustering. HBC and YS_Mac1 are highlighted in blue. (D) Violin plot of placental scRNAseq cluster enrichment scores for primitive macrophages from a Carnegie stage 10 (CS10) embryo(Zeng et al., 2019). (E) Violin plots of *HLA-DRB1* and *FOLR2* log-normalised gene expression in HBC, PAMM, YS_Mac1 and CS10 macrophages. (F) Representative flow cytometric plot and quantification of Ki67 expression by HBC (n=8). (G) Representative immunohistochemistry analysis of Ki67 expression in placental tissue sections. Black arrowheads indicate Ki67^+^cells. Scale bar = 100μm. (H) Incorporation of 5-ethynyl-2’-deoxyuridine (EDU) into FACS- isolated HBC after 18 hour culture, with and without the addition of M-CSF (n= ≥4), *p*-value calculated by one-way ANOVA. (I) UMAP visualisation of 1,091 HBC single cell transcriptomes identifying two proliferating HBC populations. (J) Dotplot heatmap of log-normalised gene expression of genes associated with stages of the cell cycle in HBC clusters. Dot size represents fraction of cells with non-zero expression. (K) UMAP visualisation of HBC with cells coloured by predicted cell-cycle state, as determined by cell-cycle scoring, with RNA velocity vector field projection calculated from all genes in all cells (black arrows) overlain. Data are represented as mean ±SEM (F) or mean alone (H).

HBC are also highly enriched for a gene signature from YS-derived embryonic macrophages from a Carnegie stage 10 embryo (~4 weeks post conception) (CS10 mac) from an additional dataset(Zeng et al., 2019) (**Figure 2D, Supplementary File 1**). PAMM display intermediate levels of enrichment for the CS10 mac gene signature. This is likely due to conserved myeloid genes not specific to ‘primitive’ macrophages within the gene signature, as it was generated via comparison between CS10 macs and non-immune cells in that dataset (**Materials and Methods**). Analysis of individual genes reveals further similarity between HBC, YS_Mac1 and CS10 mac, on the basis of *HLA-DRB1* and *FOLR2* expression (**Figure 2E**).

Due to the transcriptional similarity between HBC and ‘primitive’ macrophages, we hypothesised that HBC would be maintained in the tissue via local proliferation. We find that ~11% of freshly isolated HBC express Ki67 by flow cytometry (**Figure 2F, S1I**) and identify Ki67^+^ cells within the stroma of placental villi by immunohistochemistry (IHC) (**Figure 2G**). Furthermore, during overnight culture, ~1.5% of FACS-isolated HBC incorporate 5-Ethynyl-2’-deoxyuridine (EDU), and incorporation is slightly elevated by the addition of macrophage colony-stimulating factor (MCSF) to the cultures (**Figure 2H**). Directed analysis of the HBC cluster within the placental scRNAseq dataset identifies 2 proliferating populations, pHBC-1 and pHBC-2 (**Figure 2I**). These clusters express genes associated with distinct stages of the cell cycle *(PTTG1, CDC20, UBE2C, CDK1, PCNA* and *MCM5)* (**Figure 2J**), and cell cycle scoring assigns pHBC-1 and pHBC-2 to the S and G2/M phases of the cell cycle respectively (**Figure 2K**). RNA velocity vectors, derived by calculating the ratio between spliced and unspliced reads of each gene within each cell(La Manno et al., 2018), demonstrate a clear path of HBC through the cell cycle (**Figure 2K**). No subpopulations are observed within non-proliferating HBC, allowing us to isolate them as a single population for functional assays.

Together these data show that HBC are transcriptionally similar to macrophage populations generated through primitive haematopoiesis and are a homogenous population, proliferating within placental villi, suggesting that they arise without a monocyte intermediate.

### PAMM are heterogeneous

Our flow cytometric analysis clearly shows that PAMM consist of 2 major populations, HLA-DR^hi/lo^FOLR2^-^ cells (PAMM1) and HLA-DR^hi^FOLR2^hi^ cells (PAMM2) (**Figure 3A**). Subsequently, directed reanalysis of PAMM within the scRNAseq dataset reveals further heterogeneity on the basis of *CD9,* but not *FOLR2* expression (**Figure 3B**). Adding CD9 to our flow cytometry panel, the HLA-DR^hi/lo^FOLR2^-^ (PAMM1) cells are split into two populations (**Figure 3C**). To determine if either of these populations are circulating maternal monocytes, we added the monocyte maker CCR2 to the panel and performed flow cytometry on matched maternal blood. FOLR2^-^CD9^hi^CCR2^lo/int^ cells are not present in matched maternal blood, indicating that they are macrophages with a phenotype specific to the placental niche. The remaining PAMM1 cells do however share a similar phenotype with maternal peripheral blood monocytes (**Figure 3D**). Therefore, we subdivided PAMM1 into two populations: PAMM1a, FOLR2^-^CD9^hi^CCR2^lo/int^ (macrophages), and PAMM1b, FOLR2^-^CD9^-/int^CCR2^+^ (monocytes) (**Figure 3E, S1J**).

**Figure 3.**
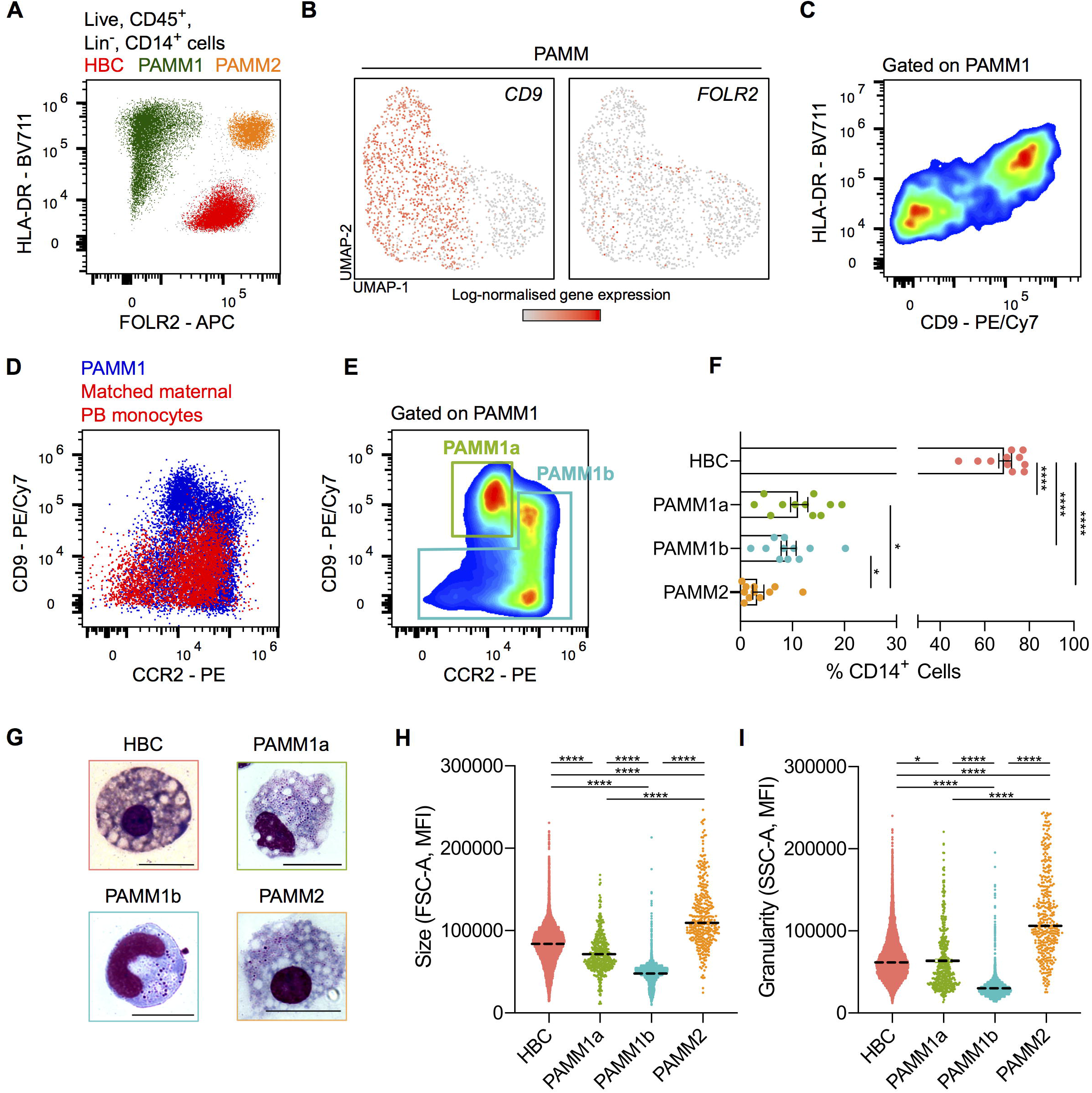
PAMM are a heterogeneous population, comprised of three subsets based on their expression of FOLR2, CD9 and CCR2 expression. (A) Expression of FOLR2 and HLA-DR by flow cytometry reveals three major populations of placental macrophages: HBC (red), PAMM1 (green) and PAMM2 (orange). (B) UMAP visualisation of 1,687 PAMM single cell transcriptomes from first trimester placenta, with overlays of *CD9* and *FOLR2* log-normalised gene expression. (C) Heterogeneous expression of CD9 within PAMM1 by flow cytometry. (D) Overlay flow cytometric plots of PAMM1 (blue) and peripheral blood (PB) monocytes from matched maternal blood (red) of CD9 and CCR2 expression. (E) Flow cytometric plot of CD9 and CCR2 expression within PAMM1, showing representative gates for the identification of PAMM1a and PAMM1b. (F) Enumeration of HBC and PAMM populations as a percentage of total CD14^+^ cells from placental cell suspensions (n=11). *p*-values were calculated by one-way ANOVA with Tukey’s multiple-comparisons test. (G) Representative Giemsa-Wright stained cytospins of HBC and PAMM subsets isolated by FACS. Scale bars = 20μm. (H) Forward scatter (FSC-A) and (I) Side scatter (SSC-A) mean fluorescence intensity (MFI) of HBC and PAMM subsets. *p*-values were calculated by one-way ANOVA with Tukey’s multiple-comparisons test. Data are represented as mean ±SEM (F) or mean alone (H, I). \**p*≤0.05, *\*\*\*\*p*≤0.0001.

HLA-DR^hi^FOLR2^hi^ cells (PAMM2) are rare in placental samples (~3% of placental CD14^+^ cells) (**Figure 3F**). Decidual macrophages also express FOLR2 and HLA-DR (**Figure S2A, S2B**) and it is likely that PAMM2 are maternal decidual macrophages that will contaminate placental samples. Although PAMM2 do not form a distinct cluster in the placental scRNAseq dataset, combined analysis of placental, decidual and maternal blood scRNAseq datasets (**Figure S2C**) reveals that HLA-DR^+^ FOLR2^+^ decidual macrophages (dMac2) (**Figure S2D)** are found in low numbers in the placental digests (**Figure S2E**)(Vento-Tormo et al., 2018). Cytospins and analysis of cell granularity and size by flow cytometry shows that HBC, PAMM1a and PAMM2 are large, granular cells with morphologies typical of macrophages - large vacuoles and pseudopods (**Figure 3G, 3H, 3I**). PAMM1b are comparatively smaller in size, and their morphology is typical of blood monocytes (**Figure 3G, 3H, 3I**). In line with their phenotypic and morphological properties, we find that PAMM1b are transcriptionally similar to adult circulating classical monocytes (**Figure S2F**). However, PAMM1b display increased expression of 150 genes, including chemokines, in comparison to maternal blood classical monocytes (**Figure S2G**). PAMM1a are not present within the decidua (**Figure S2A**), indicating their phenotype probably reflects adherence to the SCT. Of the three PAMM populations identified, PAMM1a are the most abundant, representing ~11% of the CD14^+^ cells in placental isolates (**Figure 3F**).

In conclusion PAMM can be subdivided into different populations. PAMM1b are monocytes, PAMM1a are macrophages that are specific to the placental surface, whilst PAMM2 are contaminating decidual macrophages.

### PAMM1a adhere to sites of injury on the placental surface and secrete factors involved in tissue repair

We next sought to determine the potential role of PAMM1a in healthy pregnancy. To investigate what changes in gene expression occur during the transition from PAMM1b (monocytes) to PAMM1a (macrophages) we performed Slingshot trajectory analysis(Street et al., 2018) (**Figure 4A**). Genes associated with monocyte identity and function, including *S100A8, S100A9* and *LYZ,* are downregulated along the trajectory (**Figure 4B**). Upregulated genes included macrophage markers *CD63, CD68, CD36* and *GPNMB,* and a subset of genes associated with tissue remodelling (including *LPL, MMP7,* and *MMP9)* (**Figure 4C**). The elevated surface expression of LOX-1 (the receptor encoded by *OLR1),* CD63, CD68 and CD36 by PAMM1a are verified by flow cytometry (**Figure 4D**).

**Figure 4.**
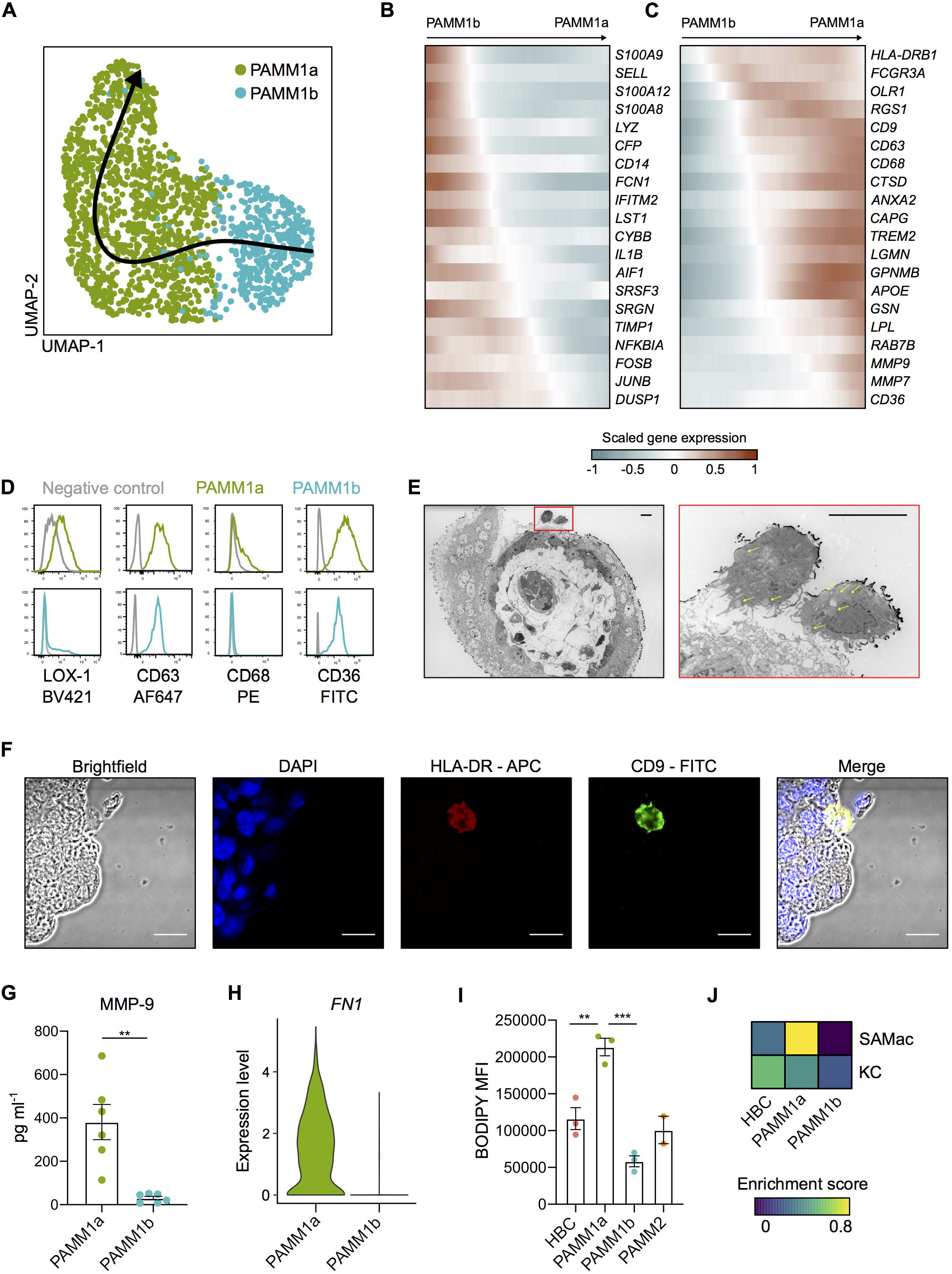
PAMM1 undergo a monocyte-to-macrophage transition and adopt a tissue-repair phenotype on the placental surface. (A) UMAP visualisation of 1,687 PAMM single cell transcriptomes with Slingshot trajectory overlain. (B) Heatmaps of smoothed scaled gene expression of selected genes which are downregulated and (C) upregulated during PAMM1b to PAMM1a differentiation, ordered according to Slingshot trajectory. (D) Relative surface expression of markers identified in (C) in PAMM1a (green) and PAMM1b (cyan), compared to FMO control (grey), measured by flow cytometry. Representative plots of n=3. (E) Transmission electron microscopy of first trimester placental villous cross-section, PAMM1a can be observed on the placental surface, localised to sites of damage to the syncytial layer (red inset). PAMM1a are loaded with lipid droplets (yellow arrows). Scale bars = 20μm. (F) Identification of CD9^+^ (green) HLA-DR^+^(red) PAMM1a cells on the surface of a 9wk EGA placental sample by fluorescence microscopy. Representative image of n=3. Cell nuclei are stained with DAPI (blue). Scale bars = 20μm. (G) Secretion of MMP-9 by FACS-isolated PAMM1a and PAMM1b after overnight culture (n=6). *p*-value calculated by unpaired t-test. (H) Log-normalised gene expression of fibronectin *(FN1)* in PAMM1a and PAMM1b clusters, as determined from scRNAseq data. (I) Analysis of intracellular neutral lipid content by flow cytometry following staining with BODIPY; mean fluorescence intensity (MFI) of HBC and PAMM subsets is shown. *p*-values calculated by one-way ANOVA with Tukey’s multiple-comparisons test. (J) Heatmap of placental macrophage mean enrichment scores for Kupffer cells (KC) and scar-associated macrophage (SAMac) gene signatures(Ramachandran et al., 2019). Data are represented as mean ±SEM. \*\**p*≤0.01, \*\*\**p*≤0.001.

Breaks occur in the SCT *in vivo* in healthy pregnancies(Burton and Watson, 1997). Fibrin deposits together with macrophages are characteristically seen at the sites of syncytial damage(Pierleoni et al., 2003; Burton and Watson, 1997). We identify PAMM1a adhered to sites of damage on the SCT by electron (**Figure 4E, S2H**) and fluorescent microscopy (**Figure 4F**). PAMM1a secrete matrix metalloproteinase (MMP)-9 (detected by Luminex assay after FACS isolation and overnight culture) (**Figure 4G**) and strongly express fibronectin (mRNA) (**Figure 4H**). Transmission electron microscopy reveals PAMM1a are laden with lipid droplet-like structures (yellow arrows **Figure 4E**). Staining with BODIPY, a dye that specifically labels neutral lipids, confirms that PAMM1a are highly loaded with lipid droplets (**Figure 4I, S2I, S2J**). Lipid droplet formation in macrophages can be induced by the uptake of apoptotic cells(Ward et al., 2018; Ford et al., 2019; D’Avila et al., 2011), suggesting PAMM1a function in the clearance of cellular debris and repair of the SCT following damage. If this is the case, PAMM1a might display transcriptomic similarities to macrophages in damaged, fibrotic tissues. Indeed, we find that PAMM1a, but not HBC, are strongly enriched for a gene signature from a population of scar-associated macrophages found in human cirrhotic livers(Ramachandran et al., 2019)(**Figure 4J, Supplementary File 1**).

To summarise, we have identified PAMM1a on the SCT, and these cells are likely to function in essential repair of the placental barrier.

### Hofbauer cells produce factors that promote placental angiogenesis

HBC, PAMM1a and PAMM1b populations were isolated by FACS from placental digests, cultured overnight and their secretion of cytokines and growth factors was determined by Luminex (PAMM2 cell yields were too low for functional assays) (**Figure 5A, S3A**). The secretion profile of PAMM1a and PAMM1b differs substantially from HBC reflecting their maternal origin. PAMM1b secrete increased amounts of the proinflammatory cytokines IL-1β and IL-6 in comparison with PAMM1a (**Figure 5A**), consistent with a monocyte to macrophage transition. In comparison with PAMM1a and PAMM1b, HBC secrete both VEGF-A and low levels of FGF2, growth factors involved in placental growth and angiogenesis(Burton et al., 2009a; Arany and Hill, 1998) as well as high levels of osteopontin (OPN), that has a role in implantation and placentation(Johnson et al., 2003). Surprisingly, HBC also secrete factors that are typically associated with inflammation such as IL-8, CCL-2, 3 and 4. However, these factors also have pro-angiogenic properties, a more likely role in the context of the placenta(Shi and Wei, 2016; Lien et al., 2018; Wu et al., 2008; Salcedo et al., 2000; Stamatovic et al., 2006). HBC expressed tissue inhibitor of metalloproteinase (TIMP)-1 and MMP-9, both of which are involved in remodelling of placental vessels(Luizon et al., 2014; Plaks et al., 2013).

**Figure 5.**
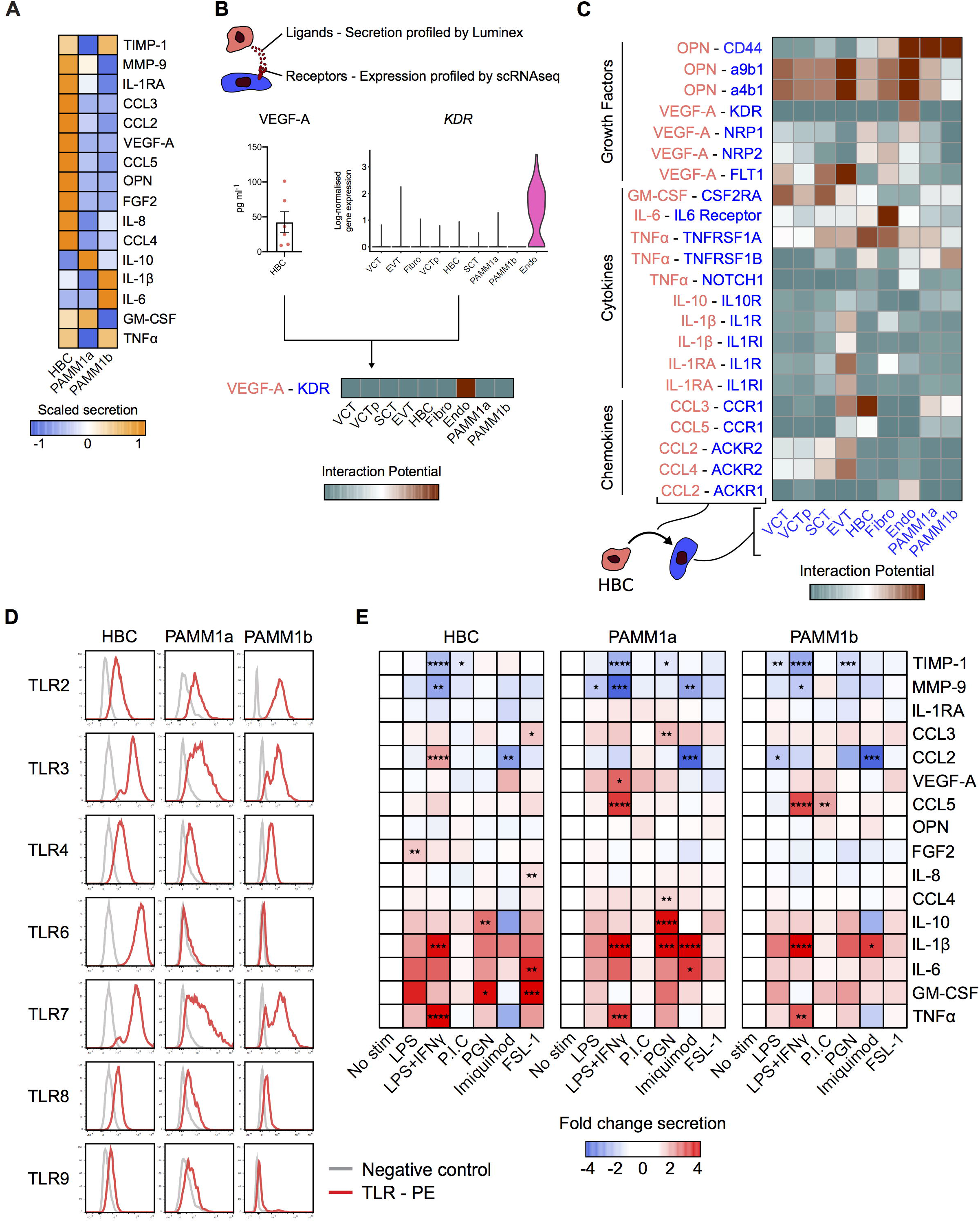
HBC and PAMM subsets display distinct cytokine secretion profiles at the steady state and in response to TLR stimulation. (A) Heatmap of average scaled cytokine, chemokine and growth factor secretion from FACS-isolated HBC, PAMM1a and PAMM1b after overnight culture without stimulation (n=6). (B) Schematic representation of inferred cell-cell interactions from Luminex and scRNAseq data. Lower panel shows an example of predicted interactions between HBC and other placental cells based on VEGF-A and KDR. (C) Heatmap of predicted interactions between HBC (red) and other placental cell populations (blue). Interaction potentials were calculated from expression of ligands determined by protein secretion, and scRNAseq expression of cognate receptors. (D) Relative flow cytometric expression of TLRs in HBC, PAMM1a and PAMM1b compared to FMO control (grey). Plots are representative of n=3. (E) Heatmaps showing the fold-change in cytokine secretion of FACS-isolated HBC, PAMM1a and PAMM1b cultured overnight with TLR stimulation, relative to no stimulation (n=6). *p*-values were calculated by two-way ANOVA with Dunnett’s multiple-comparisons test. \**p*≤0.05, \*\**p*≤0.01, \*\*\**p*≤0.001, \*\*\*\**p*≤0.0001.

To determine which cells respond to factors secreted by HBC, we generated a measure of the interaction potential between HBC and other placental cells by combining Luminex protein secretion data with scRNAseq gene expression data for cognate receptors (**Figure 5B**). Our analysis reveals predicted targets of HBC signalling (**Figure 5C, S3B**). Endothelial cells are the main target of VEGF-A secretion, mediated by the expression of kinase insert domain receptor (KDR) and neuropilin 1 (NRP1). OPN is also predicted to signal to endothelial cells, via CD44 and integrin complexes, interactions which are known to promote angiogenesis(Dai et al., 2009; Poggio et al., 2011). HBC-endothelial cell interactions are also facilitated by their close proximity within placental villi (**Figure S3C**). Additionally, HBC are predicted to signal to placental fibroblasts via IL-6, and to villous cytotrophoblast via both OPN and granulocyte-macrophage colony-stimulating factor (GM-CSF).

In summary, we have identified factors that HBC secrete that are likely to promote placental growth and homeostasis through interactions with endothelial cells, fibroblasts and trophoblast.

### Hofbauer cells are responsive to TLR stimulation

The placenta is a crucial barrier protecting the fetus from vertical infections, and HBC are the only fetal myeloid cells in the first trimester placenta. However, their role in defending the fetus from infection remains unclear. In addition, whether ‘primitive’ macrophages have the capacity to detect and respond to microbial stimuli is unknown. We therefore next asked whether HBC are responsive to Toll-like receptor (TLR) stimulation. TLRs drive specific immune responses through the recognition of distinct pathogen-associated molecular patterns, derived from a range of microbes(Kawasaki and Kawai, 2014).

The TLR expression profile of HBC analysed by flow cytometry is distinct from PAMM populations (**Figure 5D, S3D**); while HBC express TLR2, 3, 4, 7 and 8, their expression of TLR-6 is elevated in comparison with PAMM1a and 1b. TLR9 expression is low to negative in HBC, PAMM1a and PAMM1b. Interestingly, TLR expression is poorly captured by scRNAseq (**Figure S3E**), highlighting potential issues with over-reliance on gene expression data alone.

We next determined the response of HBC, PAMM1a and PAMM1b to TLR stimulation by analysing their production of cytokines and growth factors after overnight stimulation with TLR agonists. Due to differences between subsets in their baseline expression of cytokines and growth factors, as demonstrated in **Figure 5A**, expression levels are normalised to unstimulated controls (**Figure 5E, S4**). The response of HBC to TLR stimulation is specific to the agonist used, with LPS+IFNg and FSL-1, a TLR 6 agonist, having the greatest impact. A combination of LPS and IFNg impairs the ability of HBC to produce factors important in tissue remodelling, including TIMP-1 and MMP-9, and increases the secretion of IL-1β and TNFa. FSL-1 increases HBC production of CCL3, IL-8, IL-6 and GM-CSF. In contrast, PAMM1a and PAMM1b did not respond to FSL-1. These data show that ‘primitive’ HBC are capable of recognising and responding to microbial stimulation and highlight the distinct responses of HBC compared to maternal PAMM1a and PAMM1b.

### Hofbauer cells demonstrate microbicidal capacity

Given that breaks in SCT could provide a placental entry point for microbes and HBC are responsive to TLR stimulation, we next sought to determine if HBC have the mechanisms in place to kill microbes. Although mechanisms utilised by adult macrophages to kill microbes are well described in the literature, it remains unclear if ‘primitive’ macrophages such as HBC can utilise these.

HBC highly express receptors involved in phagocytosis, including CD64 (binds to IgG immune complexes), the mannose receptor CD206 (**Figure 1G**), and the scavenger receptors CD163, AXL and TIM1 (recognises phosphatidylserine (PS) and is critical for the uptake of apoptotic cells(Kobayashi et al., 2007))(**Figure S5A**). In line with these findings HBC display increased phagocytic capacity of YG beads (**Figure 6A**) and CFSE-labelled *Escherichia coli* (**Figure S5B**) in comparison with PAMM1a. Cells cultured at 4°C and in the presence of cytochalasin D (an inhibitor of actin polymerisation) were used as controls for beads bound to the cellular surface.

**Figure 6.**
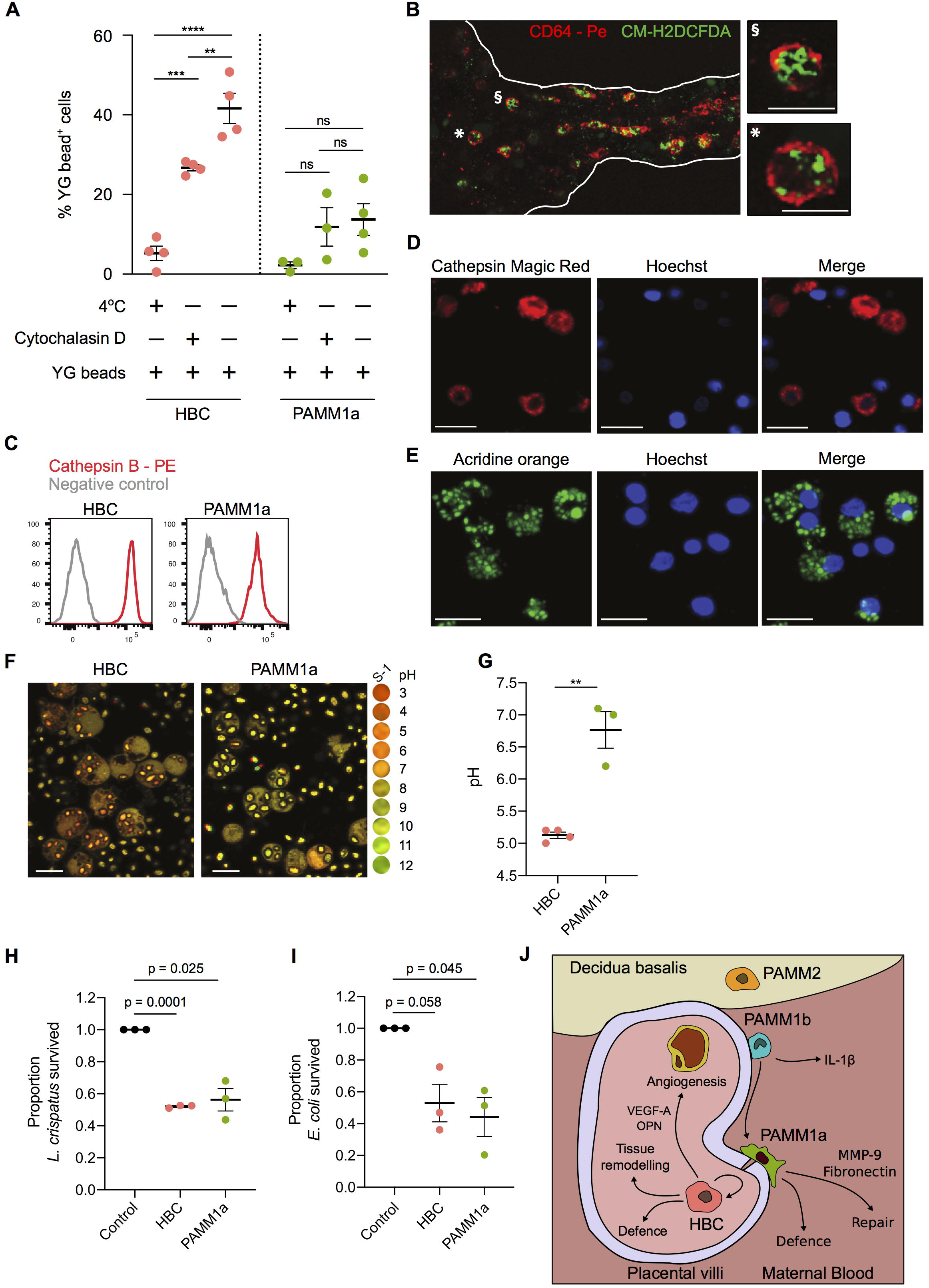
HBC are capable of mounting a microbicidal response. (A) Phagocytosis of YG beads by FACS-isolated HBC and PAMM1a measured by flow cytometry. *p*-values were calculated by two-way ANOVA with Tukey’s multiple-comparisons test. (B) Whole-mount immunofluorescence microscopy of a placental villus showing CD64 expression (red) and ROS-dependant probe CM-H2DCFDA (green). Edges of villus are indicated by white lines. Right panels, magnification of individual cells, denoted by symbols. Scale bars = 20μm. (C) Relative expression of cathepsin B in HBC and PAMM1a to FMO control (grey), measured by flow cytometry. (D) Cathepsin B activity in FACS-isolated HBC, co-cultured with zymosan particles, determined by cathepsin B Magic Red™ staining. Scale bars = 20μm. (E) Acridine orange staining of lysosomes in FACS-isolated HBC, co-cultured with zymosan particles. Scale bars = 20μm. (F, G) Comparison of the phagosomal pH of HBC and PAMM1a. (F) Representative images of phagosomal pH of FACS-isolated HBC and PAMM1a after coculture with Carboxy SNARF ®-1 -labelled zymosan particles for 20 minutes. The cytosol is labelled with 5-(and-6)-carboxy S-1 acetoxymethyl (S-1-AM) ester, a cell-permeant pH indicator. Right panel, pH scale. (G) Quantification of phagosomal pH, each data point represents an average of >100 measurements per separate donor. n≥3. *p*-value was calculated by unpaired t-test. (H, I) Rates of *Lactobacillus crispatus* (H) and *Escherichia coli* (I) killing by HBC and PAMM1a after 1 hour co-culture at a MOI of 1, relative to negative control, where no macrophages were added. *p*-values were calculated by one sample t-test. Each data point indicates a separate donor. (J) Schematic depicting locations and subset-specific roles of placental macrophages. Data are represented as mean ±SEM. ^ns^*p*> 0.05, \*\**p*≤0.01, \*\*\**p*≤0.001, \*\*\*\**p*≤0.0001.

During phagocytosis phagosomes fuse with lysosomes, resulting in production of reactive oxygen species (ROS) and protease activation. We tested if HBC can produce ROS using the ROS indicator CM-H2DCFDA. Isolated HBC make ROS even without phorbol 12-myristate 13-acetate (PMA) stimulation (**Figure S5C)**, possibly due to the stress of the cell isolation protocol. Using *ex-vivo* whole-mount immunofluorescence microscopy on placental explants, we find that the CD64^+^ HBC in the villous stroma produce ROS, indicated by CM-H2DCFCA staining (**Figure 6B**).

Both HBC and PAMM also express high levels of the protease cathepsin B (**Figure 6C**). Cathepsin B is active within HBC and PAMM1a, as demonstrated using cathepsin Magic Red™ (**Figure 6D, Figure S5D**). HBC and PAMM1a also contain lysosomal structures that were identified by the addition of acridine orange (AO) (**Figure 6E, Figure S5D**). An acidic environment in phagosomes directly aids in bacterial killing and is also important for the activation of pH-sensitive antimicrobial enzymes(Sedlyarov et al., 2018; Flannagan et al., 2015). To determine if the HBC phagosome becomes acidic during maturation, we profiled the uptake of zymosan particles tagged with a pH sensitive probe (Carboxy SNARF ®-1)(Foote et al., 2017, 2019). After allowing phagocytosis for 20 minutes, we find that the HBC phagosome becomes rapidly acidic, reaching a pH of ~4.5(**Figure 6F, 6G**). In contrast, the phagosomes of PAMM1a are more alkaline, with a pH of ~7.4 (**Figure 6F, 6G**). The pH of the PAMM1a phagosome is characteristic of antigen-presenting cells that are processing peptides for presentation(Savina et al., 2006).

Finally, to confirm that first trimester HBC have microbicidal capacity we cultured HBC with *Lactobacillus crispatus* (a microbe reported to be found in very low abundance within the 2^nd^ trimester fetal intestine (Rackaityte et al., 2020)) and *Escherichia coli* (not found in the fetus(Rackaityte et al., 2020)). We find that HBC are as efficient as PAMM1a at killing both *L. crispatus* and *E. coli,* when cultured at a MOI of 1 (**Figure 6H, 6I**) and 10 (**Figure S5E, S5F**).

Together, these data demonstrate that HBC, a population of ‘primitive’ macrophages, exhibit a range of microbicidal tools and have the capacity to kill bacteria *in vitro.*

## Discussion

Here we describe methods for the identification, isolation and characterisation of first trimester HBC. We have summarised our key findings and conclusions in **Figure 6J**. Through the application of multi-parameter flow cytometry, anti-HLA antibodies and analysis of publicly available scRNAseq datasets, we find that CD14^+^ cells obtained from first trimester placental digests contain both fetal macrophages and maternal monocytes/macrophages. Our results indicate that all previous findings on HBC from placental digests will include these maternal myeloid cells. PAMM constitute ~20-40% of isolated CD14^+^ cells and consist of 3 populations, PAMM1a, PAMM1b and PAMM2. PAMM1a are maternal monocyte-derived macrophages that have adopted a phenotype specific to the placental niche. PAMM1b are very similar to maternal monocytes, but display elevated expression of 150 genes when compared to matched maternal blood monocytes. This may reflect an adaptation to their location in the intervillous space, a unique microenvironment which can attract specific immune cells during gestation(Solders et al., 2019). However, these differences in gene expression could also reflect the isolation process for placental cells. Given their similar phenotype, PAMM2 are likely to be decidual macrophages, previously termed dMac2(Vento-Tormo et al., 2018), that contaminate the cell isolates from uterine/placental tissues from early pregnancy. They are relatively rare cells and their properties were not studied further.

Our data demonstrates that HBC are a homogenous population that are transcriptionally similar to primitive YS macrophages, further emphasising their origin through primitive haematopoiesis. While fate mapping studies using murine models have determined the origins of many tissue macrophage populations(Ginhoux and Guilliams, 2016), the origin of HBC has yet to be elucidated. The placenta is a known site of primitive haematopoiesis(Van Handel et al., 2010), but whether it occurs independently of the yolk sac or if yolk sac macrophages migrate to the placenta giving rise to HBC is unknown. HBC, YS macrophages and YS-derived macrophages from a Carnegie stage 10 embryo do not express HLA-DR. This suggests that the lack of HLA-DR is an intrinsic property of primitive macrophages, and could be used to distinguish macrophages derived from primitive and definitive haematopoiesis in other fetal tissues.

Previous studies have yielded variable findings concerning the phenotype, cytokine secretion and functions of HBC(Johnson and Chakraborty, 2012; Young et al., 2015; Pavlov et al., 2020; Loegl et al., 2016; Schliefsteiner et al., 2017; Swieboda et al., 2020), probably due to the failure to account for PAMM contamination which we show is present in placental digests. Here, using our gating strategy for the isolation of placental myeloid cell populations we find that steady-state HBC secrete a range of factors that play a role in vascularisation and the remodelling of blood vessels, such as VEGF-A, OPN, MMP-9 and TIMP-1 (Johnson et al., 2003; Dai et al., 2009; Poggio et al., 2011; Luizon et al., 2014; Plaks et al., 2013). HBC also secrete factors that are typically associated with inflammation, including IL-8, CCL-2, CCL-3 and CCL-4. IL-8 is a potent neutrophil chemoattractant(Hammond et al., 1995) but neutrophils are absent from the healthy placenta. In the context of the placenta, it is therefore likely that these factors are pro-angiogenic. For example, *in vitro* assays(Shi and Wei, 2016), using a physiological range of IL-8 (0.2 – 1 ng ml^-1^) (we found HBC produce ~128 ng ml^-1^/10^4^ cells of IL-8), have shown that IL-8 promotes the migration and canalization of human umbilical vein endothelial cells (HUVECs) and their production of VEGF-A(Shi and Wei, 2016).

The SCT layer that covers the placental surface always contains sites of damage during healthy pregnancy(Costa et al., 2017; Burton and Watson, 1997), particularly seen at bridges between 2 villi(Burton and Watson, 1997). Fibrin deposits are typically present at the sites of breaks in the syncytium(Burton and Watson, 1997). We suggest that PAMM1a are the macrophages that have been identified at sites of damage at the syncytium(Burton and Watson, 1997) and are mediators of the repair process, as they adopt a tissue-repair phenotype, and are transcriptionally similar to scar-associated macrophages in human liver fibrosis. PAMM1a are also laden with lipid droplets and are deficient in phagocytosis. This finding is resonant with data showing microglia laden with lipid droplets also display impaired phagocytosis(Marschallinger et al., 2020), although the responsible mechanisms are still unknown. PAMM1a may have reached a point of ‘saturation’ through the uptake of cellular debris at the placental surface, reducing further phagocytosis. In future studies, it will be interesting to determine the role of PAMM1a in pregnancy disorders including preeclampsia and transplacental infection.

Our work shows that while HBC are ‘primitive’ macrophages in terms of origin, they are not primitive in function, demonstrated by their response to TLR stimulation and their microbicidal capacity. The distinct response of HBC to TLR stimulation in comparison with PAMM1a reflects their TLR expression profile._For example, HBC highly express TLR-6 (binds to bacterial lipoproteins) and strongly respond to TLR-6 stimulation in comparison with PAMM1a. The elevated expression of TLR-6 by HBC is surprising given its expression is restricted to a select number of human tissues, such as the spleen(Fagerberg et al., 2014). While HBC demonstrate increased phagocytic capacity and adopt a more acidic phagosome in comparison with PAMM1a, PAMM1a are as efficient as HBC at killing both *E. coli* and *L. crispatus.* This equivalent microbicidal capacity can be explained by active cathepsin B activity and other anti-microbial mechanisms that PAMM1a may have. Given the microbicidal capacity of HBC, it is of interest to study their interaction with microbes that do cross the placental barrier and cause an adverse pregnancy outcome, such as *Listeria monocytogenes* and Zika virus. However, these areas of research were beyond the scope of this study.

The work presented in this study is limited to first trimester samples. Previous studies have investigated the properties of HBC across gestation(Goldstein et al., 1988; Ingman et al., 2010; Swieboda et al., 2020; Pavlov et al., 2020). However, studies that used placental digests have not considered contamination with PAMM populations and so interpretation of some of their findings is difficult. Using the methods described here for the isolation and study of HBC, an area of interesting future research will be to investigate how HBC phenotype and functions change throughout pregnancy.

In summary, we have provided a gating strategy that allows the study of human HBC. We have inferred the roles of these cells at the steady state and demonstrated the microbicidal capacity of human ‘primitive’ macrophages. This study adds to our understanding of human developmental immunology and provides an important framework for the field of placental biology. Future studies will now aim to determine the roles of ‘primitive’ HBC in health and disease.

## Methods and Materials

### Patient samples

All tissue samples used were obtained with written consent from participants. Decidual and placental tissues were obtained from healthy women with apparently normal pregnancies undergoing elective first trimester terminations (6-12 weeks EGA) (n=20). Peripheral blood was taken from women undergoing elective first trimester terminations (6–12 EGA). The EGA of the samples was determined from the last menstrual period. All samples were obtained with written informed consent from participants under ethical approval which was obtained from the Cambridge Research Ethics committee (study 04/Q0108/23).

### Tissue processing

Placental samples were processed immediately upon receipt. Samples were washed in PBS for 10 minutes with a stirrer before processing. The placental villi were scraped from the chorionic membrane with a scalpel and digested with 0.2% Trypsin (Pan-Biotech)/0.02% Ethylenediaminetetraacetic acid (EDTA) (Source BioScience) at 37°C with stirring, for 7 minutes. The digested cell suspension was passed through a sterile muslin gauze, and fetal bovine serum (FBS) (Sigma-aldrich) was added to halt the digestion process. The undigested tissue left on the gauze was scraped off with a scalpel and digested in 2.5ml 1mg/ml collagenase V (Sigma-Aldrich), supplemented with 50ul of 10mg/ml DNAse I (Roche) for 20 minutes at 37°C with agitation. The digested cell suspension was passed through a sterile muslin gauze and washed through with PBS. Cell suspensions from both the trypsin and collagenase digests were pelleted, resuspended in PBS and combined. Cells were layered onto a Pancoll gradient (PAN-biotech) and spun for 20 minutes without brake at 3000 rotations per minute (rpm). The leukocyte layer was collected and washed in PBS. Decidual samples and blood were processed as described previously(Huhn et al., 2020).

### Flow cytometry

Cell suspensions were stained for viability with either 1:3000 4’,6-diamidino-2-phenylindole (DAPI) (Sigma-Aldrich) or 1:1000 Zombie Aqua (Biolegend) for 20 minutes at 4°C, and washed twice in PBS. Cells were blocked in human blocking buffer (5% human serum (Sigma-Aldrich), 1% rat serum (Sigma-Aldrich), 1% mouse serum (Sigma-Aldrich), 5% FBS and 2mM EDTA) for 15 minutes at 4°C, and were incubated with antibody cocktails for 30 minutes at 4°C. Antibodies used are listed in **Supplementary Table 1**. Cells were washed and resuspended in FACS buffer (PBS containing 2% FBS and 2mM EDTA). For intracellular staining, cells were fixed and permeabilised with BD Pharmingen™ Transcription Factor Buffer (BD bioscience), according to manufacturer’s instructions. The lineage (lin) channel in flow cytometry analyses included the markers CD3, CD19, CD20, CD66b and CD335, for the removal of contaminating maternal T cells, B cells, NK cells and granulocytes. Flow cytometry was performed using a Cytek Aurora (Cytek), or cells were purified by cell-sorting using a BD FACS Aria III (BD bioscience). All flow cytometry data was analysed using FlowJo v10.6.1 (Treestar).

### Whole mount immunofluorescence microscopy

Biopsies of placental tissue (2 cm^3^) were prepared as described previously(Wang et al., 2014). Placental villi were blocked with microscopy blocking solution (1% BSA (Sigma-Aldrich), 0.25% Triton X-100 (Sigma-Aldrich) in PBS) for 15 minutes and stained with antibodies (**Supplementary Table 1**) suspended in microscopy blocking solution in 1.5ml Eppendorf tubes for 1 hour at room temperature or overnight at 4°C. The nuclear dye Hoechst 33342 (Abcam) (diluted 1:2000 in PBS) was added for 30 minutes before imaging. Whole mounts were mounted in a chamber system (POC-R2 cell cultivation system from Pecon). Imaging was performed using a Zeiss SP8 confocal LSM 700.

### Electron Microscopy

Correlative scanning and transmission electron microscopy images of PAMM on the surface of first trimester placental villi were generated as previously described(Burton, 1986).

### Immunofluorescence of placental tissue sections

First trimester placenta villous tissue and decidual sections were prepared as described previously. Slides were placed in blocking buffer for 20 minutes at room temperature, washed in PBS and incubated overnight at 4°C with antibodies (**Supplementary Table 1**). The slides were washed twice for 5 minutes in PBS, and when necessary, incubated with secondary antibodies for 1h at room temperature and washed twice for 5 minutes in PBS. The slides were then air-dried and mounted using VECTASHIELD®Antifade Mounting Medium with DAPI (Vector Laboratories). Slides were imaged using a Zeiss SP8 confocal LSM 700 (Zeiss).

### Immunohistochemistry

Slides were prepared as described previously(Sharkey et al., 1999). Antibodies used are indicated in **Supplementary Table 1**. Slides were imaged on an EVOS M5000 microscope (Thermo Fisher Scientific).

### BODIPY staining of placental cells

Placental cells were stained for flow cytometry as described above. Cells were incubated in 2ng/ml BODIPY 493/503 (Thermo Fisher Scientific) in PBS for 20 minutes at 4°C. Cells were washed in FACS buffer and acquired on a Cytek Aurora (Cytek).

FACS-isolated PAMM1a and PAMM1b were incubated in 250ng/ml BODIPY 493/503 (Thermo Fisher Scientific) in PBS for 1 hour at 37°C, fixed in 4% paraformaldehyde solution (Sigma) and washed twice in PBS. Cytospins were prepared and mounted using VECTASHIELD^®^ Antifade Mounting Medium with DAPI (Vector Laboratories) and imaged using a Zeiss SP8 confocal LSM 700.

### 5-Ethynyl-2’-deoxyuridine (EDU) incorporation assay

FACS-purified HBC were plated at a density of 10,000 cells in 100ul of Dulbecco’s Modified Eagle Medium (DMEM)(Thermo Fisher Scientific) supplemented with 10% FBS, 2.5% Penicillin Streptomycin (Sigma-Aldrich) and 20μM L-Glutamine (Sigma-Aldrich). EDU incorporation was determined using the Click-IT^TM^ Plus EdU Alexa Fluor™ 647 Flow Cytometry Assay Kit (Thermo Fisher Scientific), according to manufacturer’s instructions. Cells were incubated for 18 hours prior to harvesting and acquisition by flow cytometry.

### Cytokine production and Toll-like receptor stimulations

FACS-purified HBC, PAMM1a and PAMM1b were plated into V-bottom 96 well plates at a density of 10,000 cells in 50μl of DMEM supplemented with 10% FBS, 2.5% Penicillin Streptomycin, 20μM L-Glutamine, and 100μM Beta 2-mercaptoethanol (Sigma-Aldrich). Cells were incubated for 18 hours without stimulus, or with the following stimuli: Lipopolysaccharide (LPS) (Invivogen) 100ng/ml(Sander *et al.,* 2017), IFNγ (Novaprotein) 1000U/ml(Sander *et al.,* 2017), Polyinosinic:polycytidylic acid (poly(I:C)) (InvivoGen) 25μg/ml(Farina *et al.,* 2004), Imiquimod (Insight Technology LTD) 20μg/ml, Peptidoglycan (PGN) (InvivoGen) 10μg/ml), Pam2CGDPKHPKSF (FSL-1) (InvivoGen) (200ng/ml). After incubation, plates were spun to remove cellular debris, and supernatants were collected and stored at −80°C.

Cell culture supernatants were tested for the presence of 16 analytes using a custom 10-plex Luminex ProcartaPlex assay (Thermo Fisher Scientific), and a custom 6-plex Luminex ProcartaPlex assay (Thermo Fisher Scientific) designed to profile the expression of: CCL2, CCL3, CCL4, CCL5, FGF-2, GM-CSF, IL-1β, IL-1RA, IL-6, IL-8, IL-10, MMP-9, Osteopontin, TIMP-1, TNF-α and VEGF-A. Samples were diluted in cell culture medium at a ratio of 1:1 for the 10-plex Luminex ProcartaPlex assay, and 1:40 for the 6-plex Luminex ProcartaPlex assay. The Luminex assays were performed according to manufacturer’s instructions, and beads were ran on a Luminex LX-200 (Luminex), using xPONENT software (Luminex). Results were visualised using Prism 8 (GraphPad) and R version 3.5.1 (The R Foundation).

### Phagocytosis assays

Microsphere phagocytosis assay: Macrophages were placed in 1.5ml Eppendorfs, at 10,000 cells in 100μl PBS and human serum opsonised Fluoresbrite™ Yellow Green Microspheres 1μm (Polysciences) were added at a concentration of 10:1 for 1 hour. Controls included cells cultured at 4°C and 37°C in the presence of 10μM cytochalasin D (Sigma-Aldrich).

E.coli phagocytosis assay: *E.coli* were grown until log-phase growth in and had reached an optical density (595nm) of 0.3. Bacteria were opsonised in heat-inactivated human serum for 30 minutes at 37°C and labelled via incubation with 10μM carboxyfluorescein succinimidyl ester (CFSE) (Biolegend) for 30 minutes at 37°C. Labelled bacteria were washed 3 times in PBS prior to use. First trimester placental cell suspensions were plated at a density of 1×10^6^/ml per well in PBS. Labelled *E.coli* were added at a MOI of 10, and cultured for 1 hour at 37°C. Control wells were incubated at 4°C and at 37°C in presence of 10μM Cytochalasin-D. Plates were centrifuged at 200g for 5 minutes to promote cell-bacteria interactions. Cells were washed 3 times in 4°C PBS, and stained for flow cytometry as described above.

### Reactive oxygen species production assays

FACS isolated HBC were plated at a density of 50,000 cells in 50μl of DMEM (Gibco) supplemented with 10% FBS (Sigma-Aldrich), 2.5% Penicillin Streptomycin, 20μM L-Glutamine. Cells were stained with 1μM CM-H2DCFDA (Thermo Fisher Scientific), and were either treated with 1x cell activation cocktail (PMA-ionomycin) (Biolegend) for 30 minutes, or incubated without stimulation. Cells were washed in PBS and data acquired as described above.

*Ex-vivo* imaging of HBC ROS production was carried out by incubating placental villi with anti-CD64 antibody conjugated to PE and 1μM CM-H2DCFDA for 15 minutes. The villi were placed in Ibidi μ-Dish 35 mm and imaged using a Zeiss SP8 confocal LSM 700 (Zeiss).

### Cathepsin B activity assay and acridine orange assay

Macrophages were seeded at 10,000 cells/well in 10μl PBS on poly-L-lysine (Sigma) coated Ibidi 4 well μ-Dish plates. Zymosan bioparticles (Thermo Fisher Scientific) were added at a concentration of 10 particles per cell. Cathepsin B activity was determined using Magic Red™ (a cell-permeable and non-cytotoxic reagent that contains a cathepsin B target sequence peptide (RR)2 linked to a red (Cresyl Violet) fluorescent probe) and lysosomes were detected with acridine orange (Biorad), according to manufactures instructions.

### Phagosomal pH assay

Placental macrophage phagosomal pH measurements were performed by adapting a method using the fluorescent-sensitive pH dye SNARF-1 (S-1) with a dual emission spectrum(Foote et al., 2017). Carboxy-S-1-succinimidyl ester (Thermo Fisher Scientific) was coupled to zymosan coated beads (Thermo Fisher Scientific) and opsonised with human serum. Macrophages were cultured at 10,000 cells/well in 10μl PBS, in poly-lysine (Sigma) coated Ibidi 4 well μ-Dish plates. 5×10^5^ S-1 labelled beads were added per well to macrophages. 50μg of Carboxy S-1-acetoxymethyl (S-1-AM) ester (Thermo Fisher Scientific) was added as a cytosolic dye (0.5 μg/ml working solution). Subsequently, cells were examined under a 63x oil-immersion objective on a Zeiss SP8 confocal LSM 700 (Zeiss), where cells were excited at 555 nm and emission was measured at 560-600 nm and 600-610 nm. Over 100 measurements were made per condition. The pH scale was generated as described previously(Foote et al., 2017).

### *Escherichia coli* and *Lactobacillus crispatus* killing assays

*E. coli* and *L. crispatus* were grown overnight in LB and Man, Rogosa and Sharpe (MRS) broth respectively. Bacteria were subcultured the following day until they had reached log-phase growth and an optical density (595nm) of 0.3. Bacteria were opsonised in heat-inactivated human serum for 30 minutes at 37°C. FACS purified placental cells were plated into 96 well plates at a density of 10,000 cells in 50μl of DMEM (Gibco) supplemented with 10% FBS (Sigma-Aldrich), 20μM L-Glutamine and 25mM Hepes (Gibco). Bacteria were added at a MOI of 1 or 10, and plates were centrifuged at 200g for 5 minutes to promote cell-bacteria interactions. After incubation for 1 hour, cells were lysed in deionised water, and serial dilutions were plated onto LB agar or MRS agar plates, for *E. coli* and *L. crispatus* experiments, respectively. Colony forming units (CFU) were counted after 24 hours for E. coli and after 48 hours for *L. crispatus.*

### Analysis of publicly available single-cell RNAseq data

Single-cell RNA sequencing (scRNAseq) data of first trimester placenta(Vento-Tormo et al., 2018) was obtained at from EMBL-EBI ArrayExpress (www.ebi.ac.uk/arrayexpress), under the experiment code E-MTAB-6701. Sequencing data from placental samples were aligned using the Cell Ranger Single-Cell Software Suite (v3.0, 10x Genomics) against the GRCh38.93 human reference genome. Downstream analysis of each sample was performed using Seurat (v3.0)(Butler et al., 2018). Cells with fewer than 500 detected genes, and more than 20% mitochondrial gene expression were removed. Samples were log-normalised and integrated following the Seurat v3 Integration workflow. Clusters were identified using the *FindNeighbours* and *FindClusters* functions in Seurat. Clusters were annotated on the basis of expression of known marker genes. Uniform Manifold Approximation and Projection (UMAP) dimensionality reduction was performed using the *RunUMAP* function in Seurat, with default parameters. Significantly differentially expressed gene (DEGs) were identified using the *FindMarkers* function, using the Wilcox rank sum test, corrected for multiple comparisons.

scRNAseq data from early human fetal immune cells was obtained from GEO under the accession code GSE133345. The dataset was analysed using Seurat, and subset to include only myeloid cells. These cells were integrated with HBC and PAMM clusters from the placenta scRNAseq dataset using the reference-based integration workflow in Seurat, using the early human fetal myeloid cell object as a reference. Pearson’s correlations between annotated clusters were calculated using the average expression per cluster of the 2000 variable genes used for integration. Cell cycle scoring of HBC was performed using the *CellCycleScoring* function in Seurat, and predicted cell cycle states were overlaid onto UMAP embeddings.

Cell Ranger output files for each sample were analysed using Velocyto(La Manno et al., 2018) (python version 0.17.17). Output loom files were merged with Seurat objects in R and RNA velocity vectors were calculated using the *RunVelocity* function from the SeuratWrappers package, and projected onto UMAP embeddings using the *show.velocity.on.embedding.cor* function from the VelocytoR package.

Comparisons of cell type similarity between datasets was performed using a random forest model in the ranger R package, as previously described(Stewart et al., 2019). PBMC scRNAseq data for comparison was downloaded from 10x Genomics (https://support.10xgenomics.com/single-cell-gene-expression/datasets). Expression matrices from both datasets were subset using the union of the highly variable features detected in each dataset. The random forest model was built using the *ranger* function on the PBMC dataset and single-cell prediction scores were generated for placental cells using the *predict* function.

Single cell gene signature enrichment scores were calculated using the *AddModuleScore* function in Seurat. Gene signatures from the cirrhotic liver scar-associated macrophages (SAMacs) and Kupffer cells (KCs)(Ramachandran et al., 2019) were generated from the analysis of scRNAseq data obtained from GEO under the accession code GSE136103. In brief, the datasets were aligned and pre-processed as described above, and subset to include only the myeloid compartment. DEGs in SAMac and KC clusters were identified, and genes with log fold change > 0.5 and adjusted p value < 0.05 were used as gene signatures. DEGs were identified between yolk sac macrophages and embryonic monocytes (Bian et al., 2020), and the top fifty genes with log fold change > 0.5 and adjusted p value < 0.05 were used as the gene signatures for each population. The gene signature from CS10 macrophages(Zeng et al., 2019) was generated from the analysis of scRNAseq data obtained from GEO under the accession code GSE135202. The data was processed as described above and subset to include endothelial and hematopoietic populations. DEGs in macrophages were identified, and the top fifty genes with log fold change > 0.5 and adjusted p value < 0.05 were used as the gene signature.

Pseudotime trajectory analysis of PAMM was performed with the Slingshot R package(Street et al., 2018), and the calculated trajectory was overlain onto the UMAP embeddings. Selected genes which varied across the slingshot trajectory were plotted as heatmaps of smoothed scaled gene expression. Smoothing was performed using the *rollmean* function in the zoo R package.

### Cell-cell interactions analyses

Predicted cell-cell ligand-receptor interactions were inferred from scRNAseq data using CellphoneDB (Efremova et al., 2020), using the online tool (www.cellphonedb.org). The minimum proportion of cells in a cluster expressing a gene was set to 10%, and the number of iterations was set to 1000. Ligand-receptor pairs were subset to include only the 16 ligands profiled in Luminex experiments. An estimate for interaction potential between cells was obtained by multiplying the log-normalised cytokine secretion of each ligand from sort-purified cell populations, by the average log-normalised gene expression of each receptor in each cluster.

## Online Supplemental Material

**Figure S1** shows the digestion protocols for obtaining single cell suspensions, HLA allotype staining for matched maternal blood and decidua, relating to Figure 1B, first trimester placental scRNAseq analysis, and the full gating strategy for the isolation of HBC and PAMM populations. **Figure S2** shows the identification and characterisation of PAMM populations. **Figure S3** shows the data for cytokine, chemokine and growth factor secretion, related to Figure 5A, predicted interactions of HBC, PAMM1a and PAMM1b with other placental cells, quantification of TLR flow cytometric data related to Figure 5D, and TLR gene expression data. **Figure S4** shows the normalised cytokine, chemokine and growth factor secretion, related to Figure 5E. **Figure S5** shows the phagocytic and microbial capacity of HBC and PAMM1a. **Supplementary file 1** presents the gene signatures used for scRNAseq enrichment analyses, relating to Figures 2A, 2D and 4J. **Supplementary Table 1** provides a list of reagents, antibodies and software used throughout the study.

## Acknowledgements

We thank the following for assistance: The Flow Cytometry Core Facility at the Department of Pathology and Core staff at the Immunophenotyping Hub at the Department of Medicine, University of Cambridge. Mike Hollinshead at the Microscopy Core at the Department of Pathology, Cambridge. Kjersti Aagaard from the Department of Obstetrics & Gynecology, Division of Maternal-Fetal Medicine, Baylor College of Medicine and Texas Children’s Hospital, Houston Texas and Menna Clatworthy from the Department of Medicine, University of Cambridge for scientific discussion. Lucy Gardner, Imogen Duncan and Ritu Rani for their help in processing placental samples. We thank all donors who participated in this study and hospital staff.

This work was supported by the Wellcome Trust, Royal Society, Centre for Trophoblast Research, and Department of Pathology, University of Cambridge, UK. N.McG, is funded by a Wellcome Sir Henry Dale and Royal Society Fellowship (grant number 204464/Z/16/Z). J.T is funded by a Wellcome Trust PhD Studentship (grant number 215226/Z/19/Z). AS is funded by the MRC (grant number: MR/P001092/1).

## Author Contributions

Conceptualisation: N.McG, J.T, A.S and A.M. Methodology: N.McG, J.T, X.Z, A.A, R.D, M.D, C.L, P.N, J.C and G.B. Formal analysis: N.McG, J.T, A.A, X.Z, R.D, M.D, C.L. and G.B. Intellectual input: N.McG, J.T, A.M, A.S, F.G, G.B, X.Z and R.S. Writing: N.McG, J.T, A.S and A.M. Visualisation: N.McG and J.T. Supervision: N.McG. All authors discussed the manuscript.

## Supplementary Data

**Figure S1. Isolation and characterisation of placental macrophage populations.** (A) Schematic representation of a digestion protocol used to isolate placental cells(Tang et al., 2011). (B, C) Flow cytometric analysis of fetal myeloid cells (HLA-A2^+^), from the same sample digested with either trypsin alone (B), or (C) trypsin and collagenase. HBC (black gate) and PAMM are identified in both steps of the digestion process. (D, E) Flow cytometric analysis of maternal peripheral blood monocytes (D) and decidual CD14^+^ cells (E), matched with the placental sample shown in Figure 1 B, C. (F) UMAP visualisation of 22,618 placental single cell transcriptomes(Vento-Tormo et al., 2018). VCT – villous cytotrophoblast, VCTp – proliferating villous cytotrophoblast, SCT – syncytiotrophoblast, EVT – Extravillous trophoblast, Fibro – Fibroblasts, Endo – Endothelial cells. (G) UMAP visualisation with overlays of *CD68* and *HLA-DRB1* log-normalised gene expression. (H) Violin plots showing log-normalised gene expression of *RSP4Y1* and *XIST,* for one male fetal donor from scRNAseq dataset. (I) Flow cytometric plots showing the gating strategy and representative Ki67 staining for HBC (n=8). (J) Representative flow cytometric plots for gating strategy used to isolate HBC and PAMM populations for phenotypic, morphological and functional analysis. For the donor shown, maternal and fetal cells are HLA-A3^+^ and HLA-A3^-^ respectively.

**Figure S2. Identification of PAMM populations.** (A) Flow cytometric analysis of decidual CD14^+^ cells. Cells with a phenotype consistent with PAMM2 (FOLR2^+^ HLA-DR^+^) (red gate) and PAMM1b (blue gate) were readily identified. Cells with a phenotype consistent with PAMM1a were low in abundance (green gate). Representative flow cytometric plots from n=3 experiments. (B) Identification of HLA-DR^+^(red) FOLR2^+^ (green) macrophages in the decidua by fluorescence microscopy. Cell nuclei are stained with DAPI (blue). Scale bar = 50μm. Representative image of n=2. (C) UMAP visualisation of 9,474 myeloid cells from placenta, decidua and maternal blood(Vento-Tormo et al., 2018). Cells are coloured and labelled by cluster identity (left panel) and tissue of origin (right panel). cDC1 – conventional type 1 dendritic cells, cDC2 – conventional type 2 dendritic cells, C mono – classical monocytes, dMac1 – decidual macrophages 1, dMac2 – decidual macrophages 2, dMono – decidual monocytes, pDC – plasmacytoid dendritic cells, pHBC/pdMac2 – proliferating HBC and dMac2, NC Mono – non-classical monocytes. (D) Violin plots showing log-normalised gene expression of *FOLR2* and *HLA-DRB1* in placental, decidual and maternal blood myeloid cells. (E) Annotation of PAMM2 (placental cells within dMac2 cluster) (blue) onto original UMAP embedding of HBC and PAMM from the placental scRNAseq dataset (Figure S1F). (F) Heatmap of transcriptional similarity between placental macrophage/monocytes cell clusters and indicated PBMC populations, as determined using a random forest classification prediction. p-HBC – proliferating HBC. (G) Scatterplot showing log-normalised gene expression of PAMM1b (x-axis) and maternal blood classical monocytes (y-axis) clusters. Red dots represent genes that are differentially expressed with an adjusted p value <0.01 (Wilcox rank sum test). (H) Scanning electron micrographs of PAMM1a on the surface of a first trimester placenta, adhering to a site of damage on a branching villus. Scale bars = 20μm. (I) Representative flow cytometric histograms of BODIPY staining within HBC and PAMM subsets, compared to unstained cells (grey). (J) Images of BODIPY staining of FACS-isolated PAMM1a and PAMM1b. Scale bars = 20μm. Representative images of n=2.

**Figure S3. HBC, PAMM1a and PAMM1b secretome analysis in the steady-state and TLR expression.** (A) Cytokine, chemokine and growth factor secretion of FACS-isolated HBC, PAMM1a and PAMM1b after 18 hours in culture without stimulation, profiled by Luminex (n=6). *p*-values calculated by one-way ANOVA with Tukey’s multiple-comparisons test. Only significant *p*-values, and *p*-values approaching significance shown. (B) Heatmap of predicted interactions between HBC, PAMM1a and PAMM1b (red) and other placental cell populations (blue). Interaction potentials were calculated from expression of ligands determined by protein secretion, and scRNAseq expression of cognate receptors. (C) Whole-mount immunofluorescence of placental villi stained for CD206 (red), and CD31 (green). Images from 2 independent donors, both 9wk EGA. Scale bar = 100μm. (D) Quantification of expression of TLRs in HBC, PAMM1a and PAMM1b, profiled by flow cytometry, n=3. (E) Dotplot heatmap of log-normalised gene expression of TLR genes in HBC and PAMM scRNAseq clusters. Dot size represents fraction of cells with non-zero expression. TLR9 was not detected in the analysis. Data are represented as mean ±SEM.

**Figure S4. HBC, PAMM1a and PAMM1b secretome analysis in response to TLR stimulation.** Normalised cytokine, chemokine and growth factor secretion of FACS-isolated HBC, PAMM1a and PAMM1b after 18 hours in culture with TLR stimulation, relative to without stimulation. HBC (red), PAMM1a (green), PAMM1b (cyan). Profiled by Luminex (n=6). Data are represented as mean ±SEM.

**Figure S5. Phagocytic and anti-bacterial capacity of HBC and PAMM1a.** (A) Flow cytometric plots of scavenger receptor expression in HBC (red) and PAMM (grey). Representative flow cytometric plots of n=3 experiments. (B) Phagocytosis of CFSE-labelled *Escherichia coli (E. coli)* by HBC, PAMM1a, PAMM1b and PAMM2 subsets measured by flow cytometry. *p*-values were calculated by two-way ANOVA with Tukey’s multiplecomparisons test. (C) Representative flow cytometric plot of CM-H2DCFDA staining in FACS-isolated HBC with no stimulation (black) and with phorbol 12-myristate 13-acetate (PMA) (red), relative to no stain (grey) Representative plot from n=3 experiments. (D) Cathepsin B activity, determined by cathepsin B Magic Red™ staining, and acridine orange staining of lysosomes in FACS-isolated PAMM1a, co-cultured with zymosan particles. Scale bars = 20μm. (E, F) Rates of *Lactobacillus crispatus (L. crispatus)* (E) *and E. coli* (F) killing by HBC and PAMM1a after 1 hour co-culture at a MOI of 10, relative to negative control, where no cells were added. *p*-values were calculated by one sample t-test. n ≥ 2. Data are represented as mean ±SEM. ^ns^*p*> 0.05, \**p*≤0.05, \*\*\*\**p*≤0.0001.

**Supplementary Table 1.**
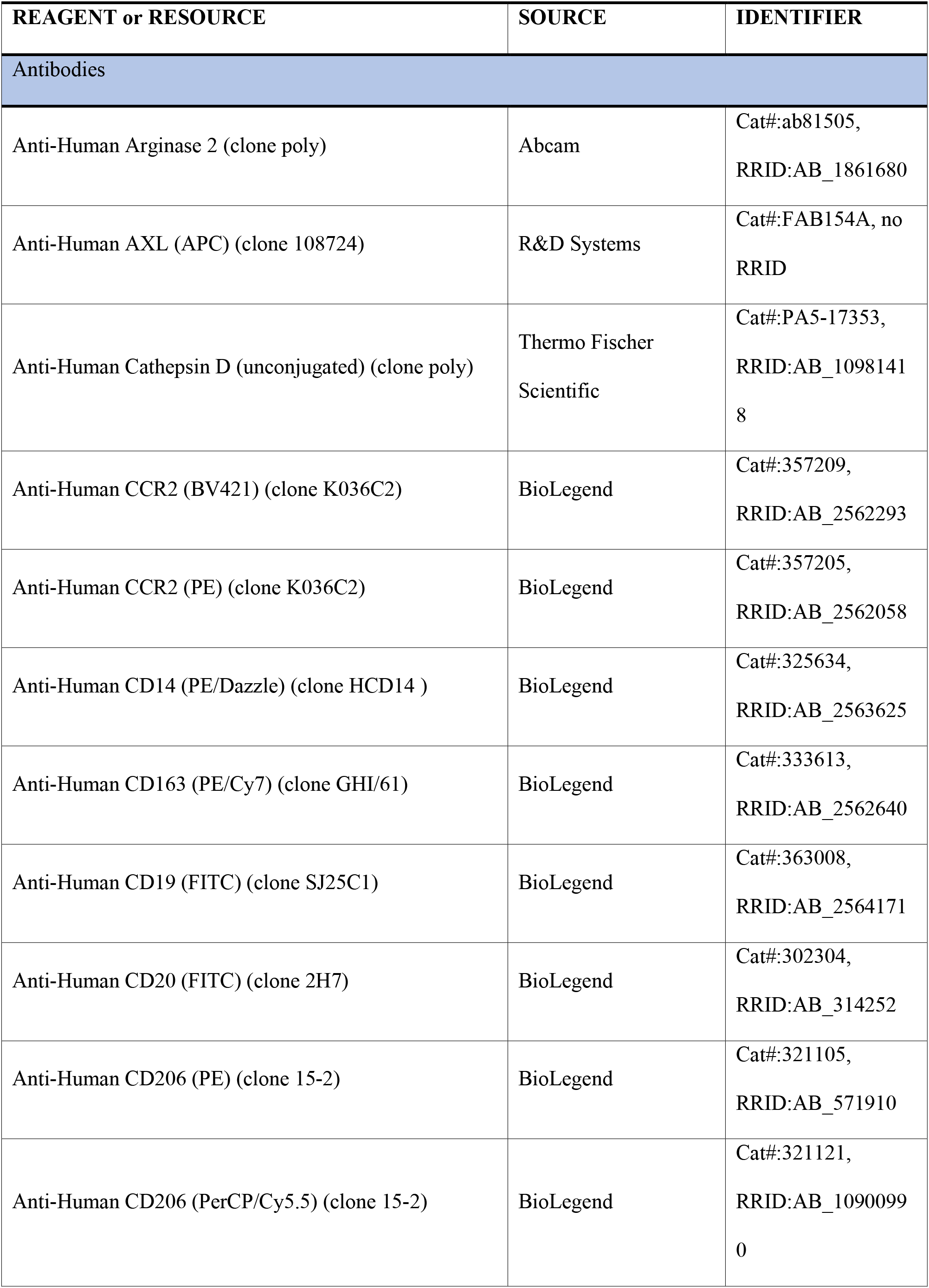
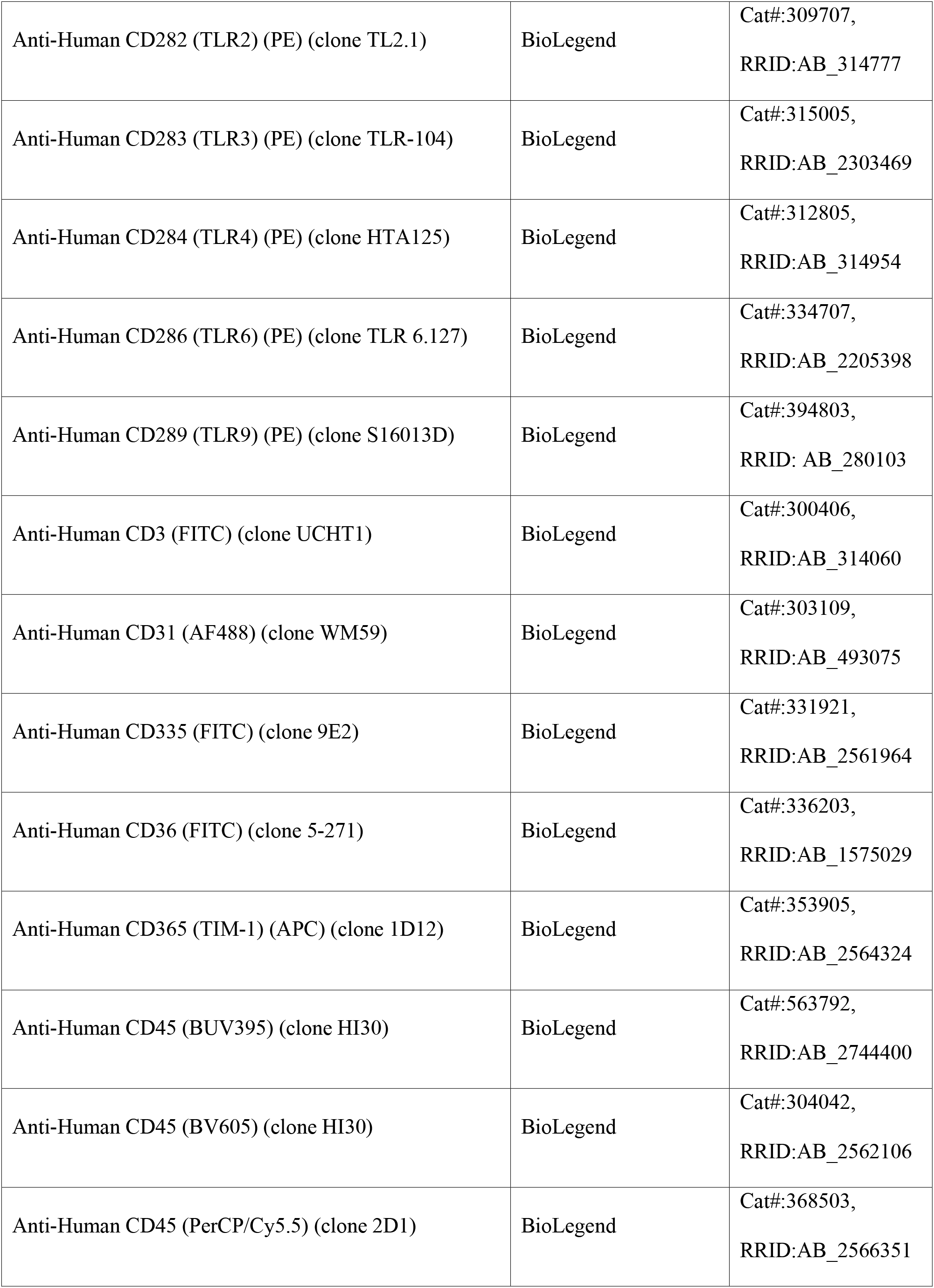
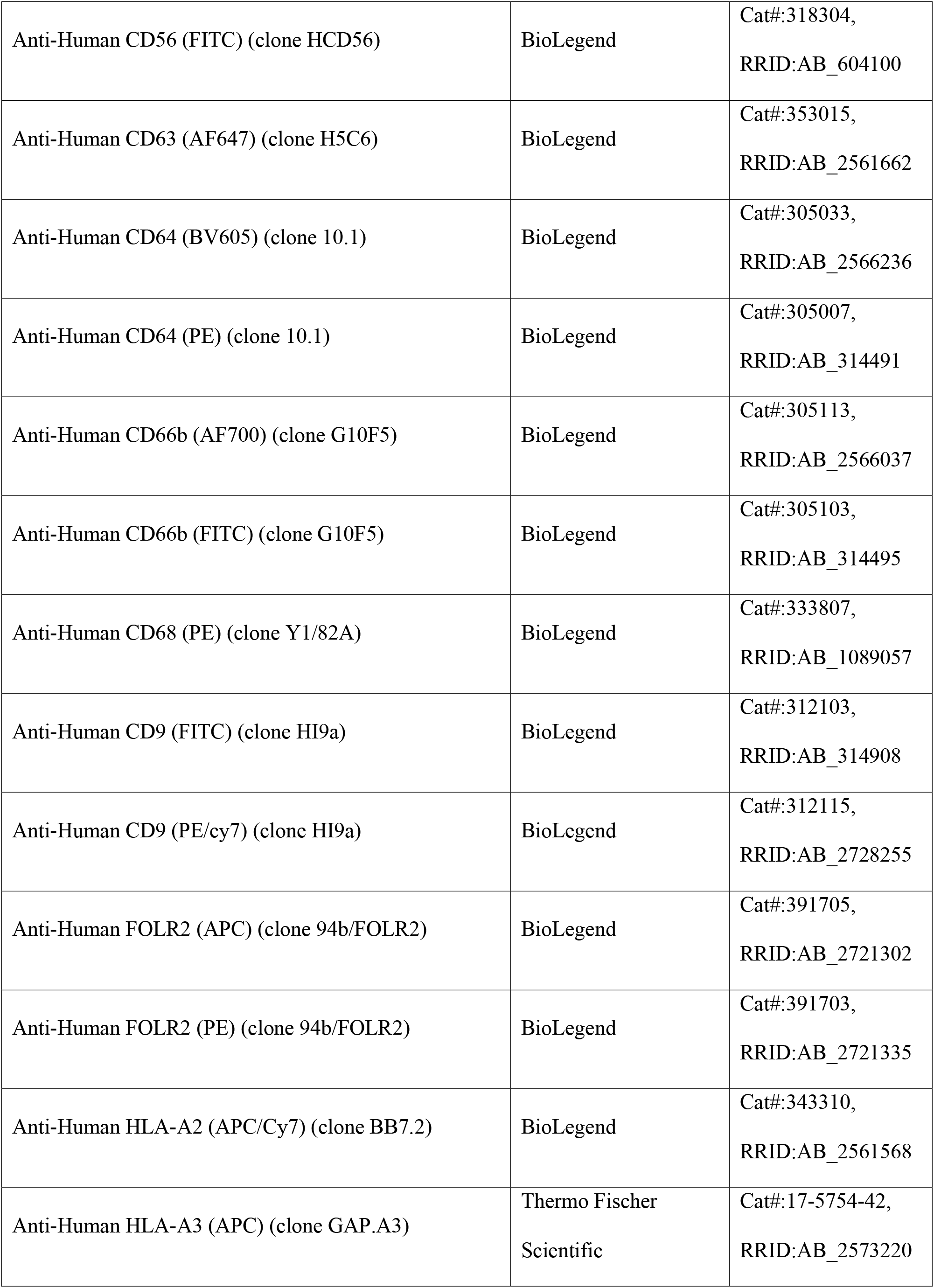
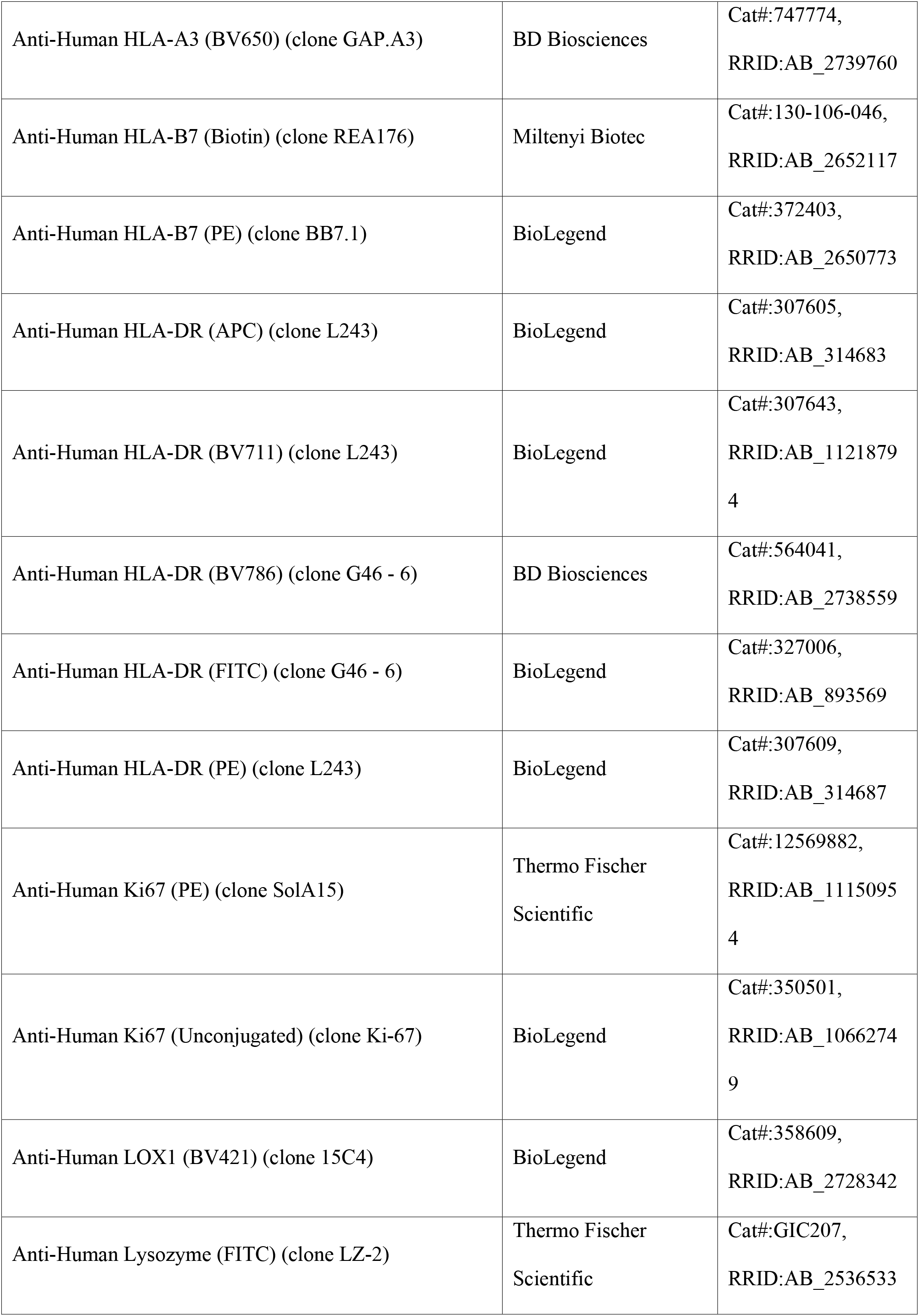
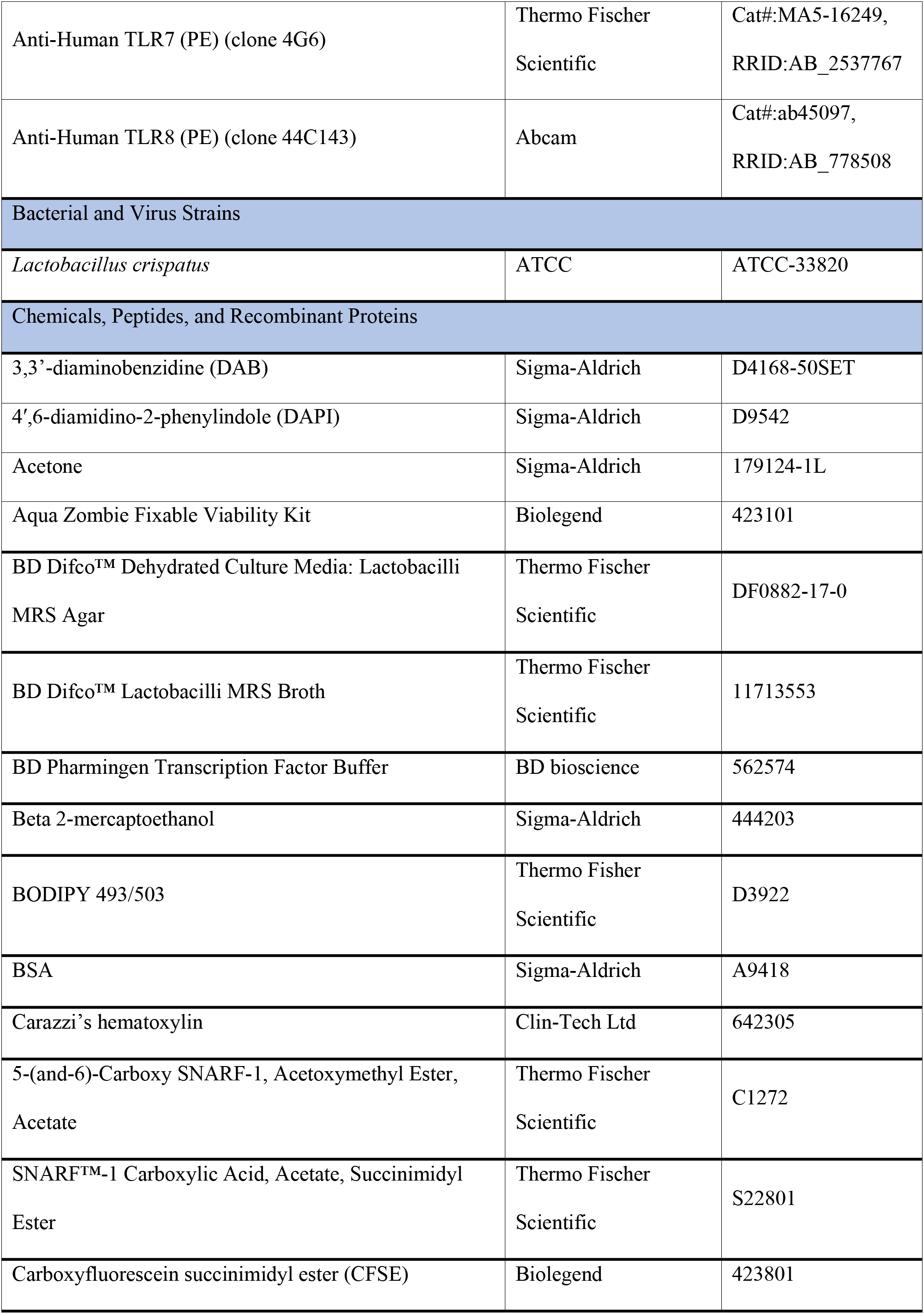
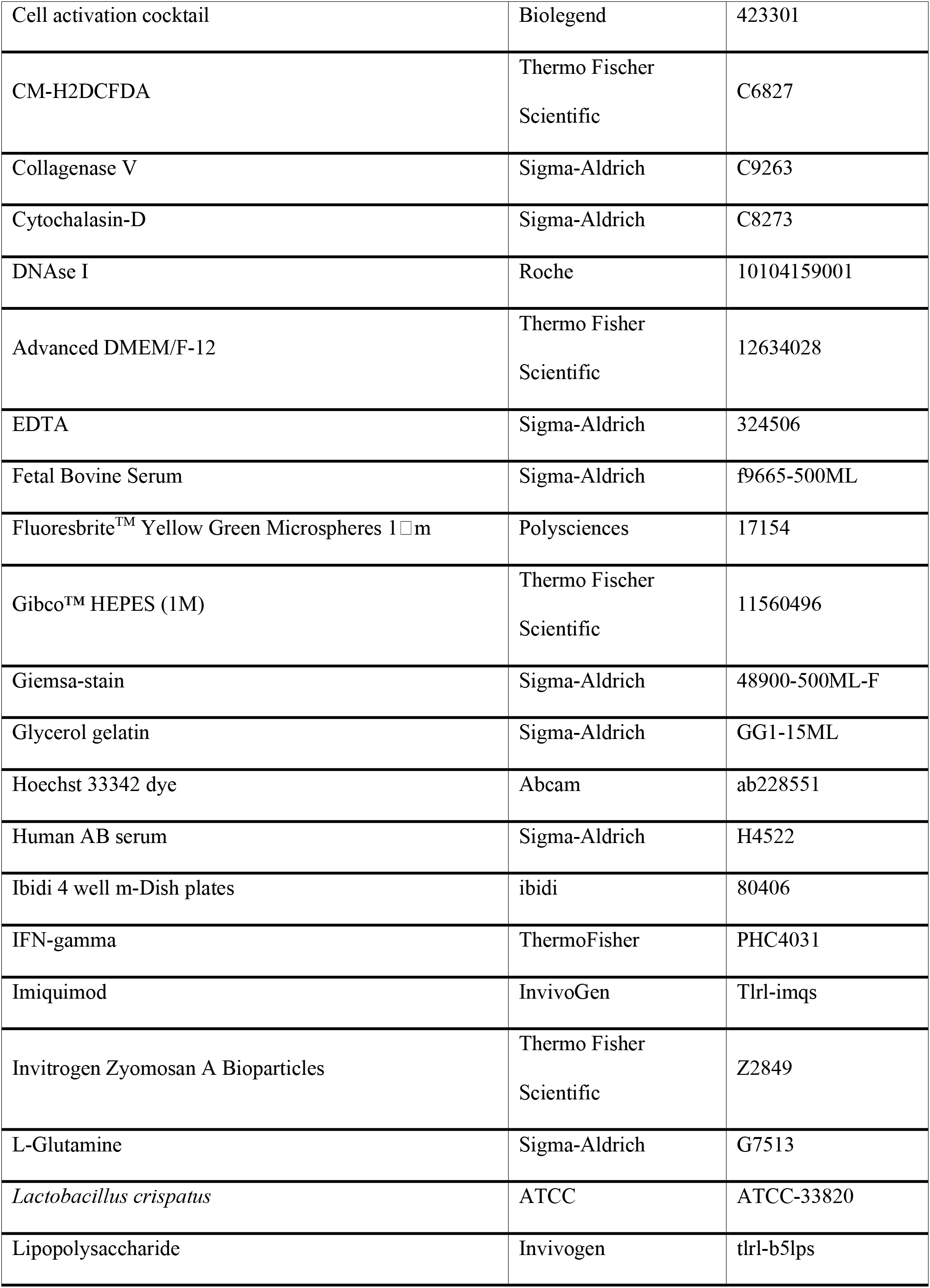
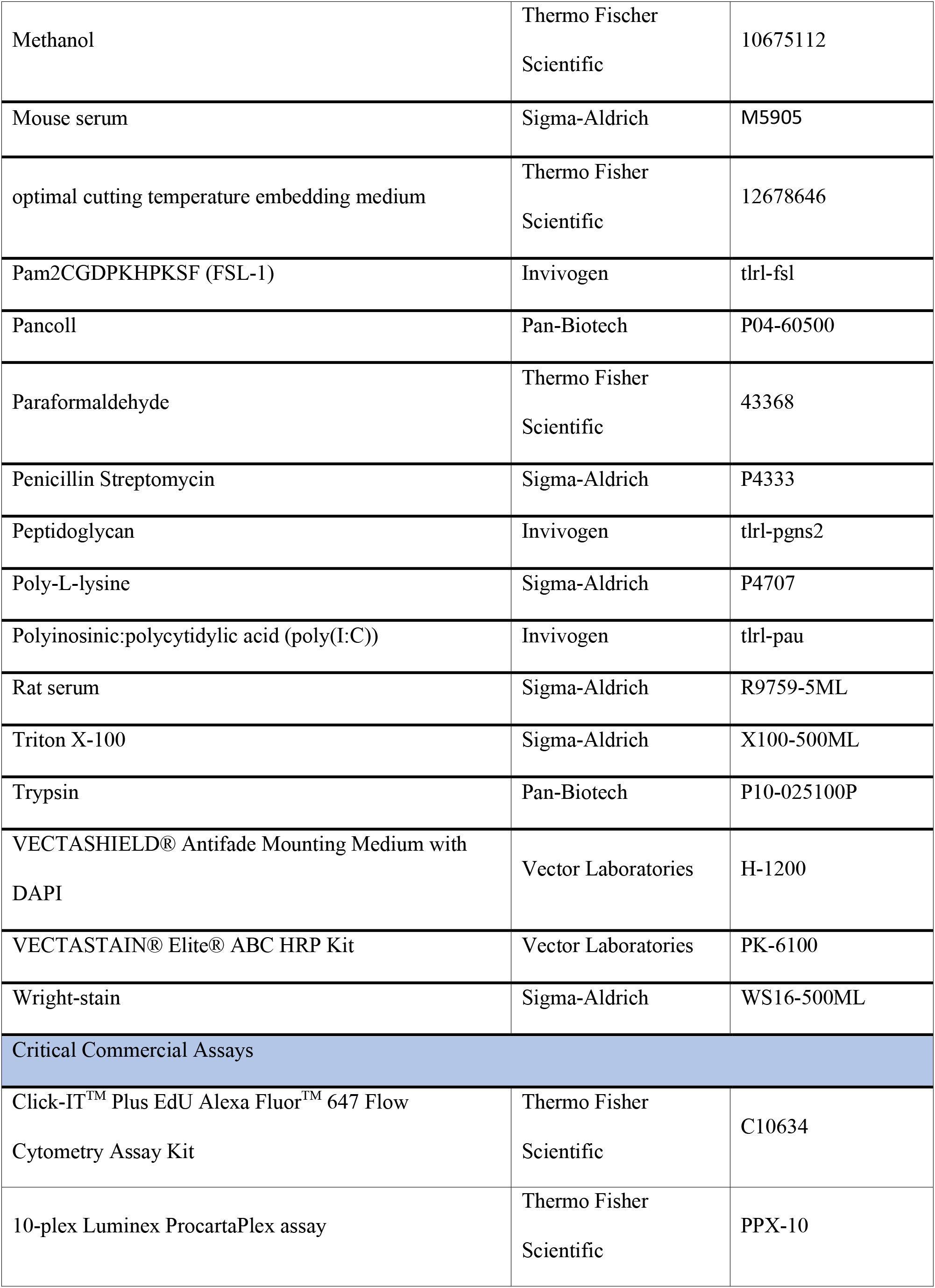
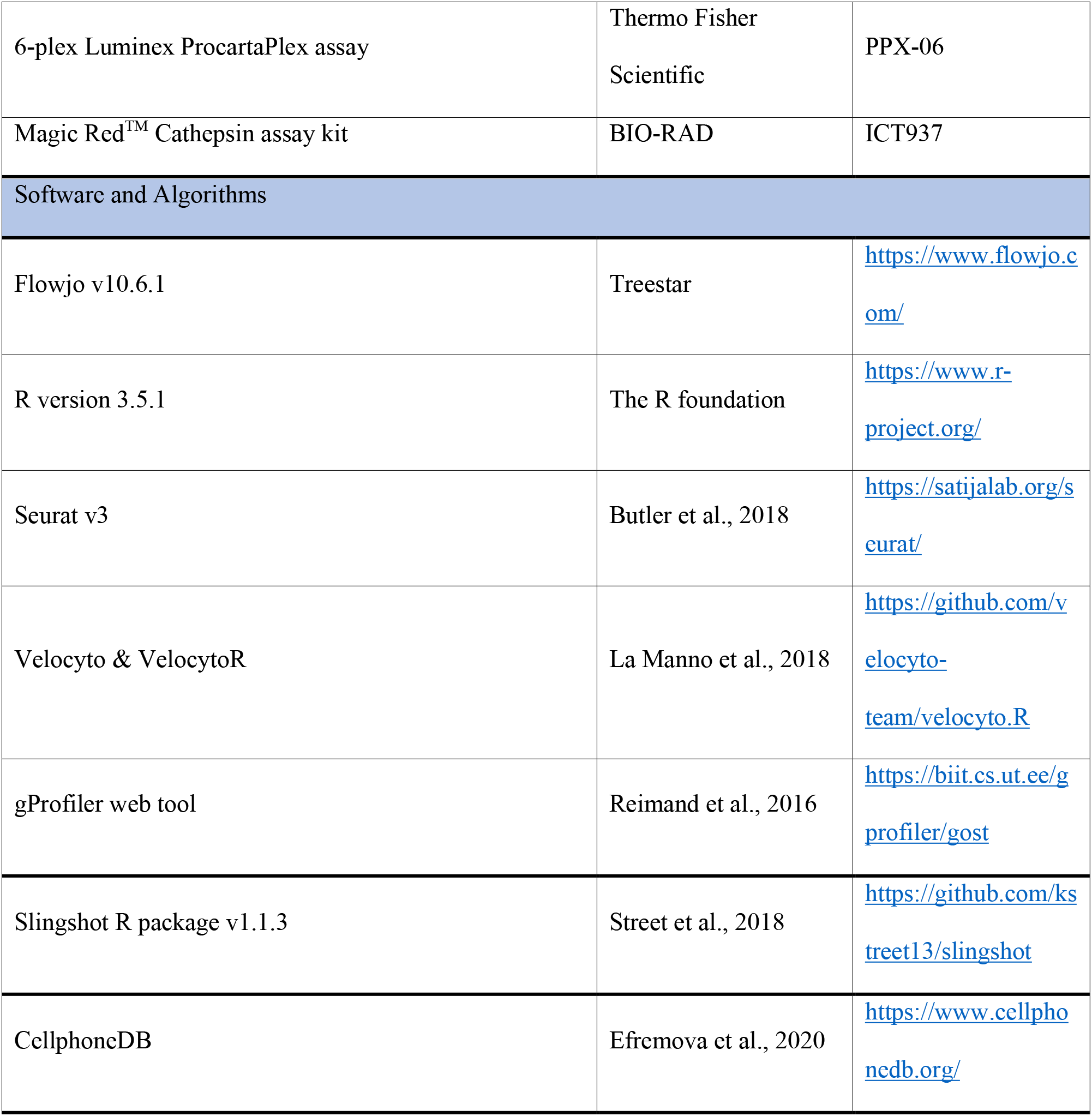
Antibodies, reagents and software used.

