## Supplementary figures and images for "Phenotypic and functional characterisation of first trimester human placental macrophages, Hofbauer cells"

### Figure S1

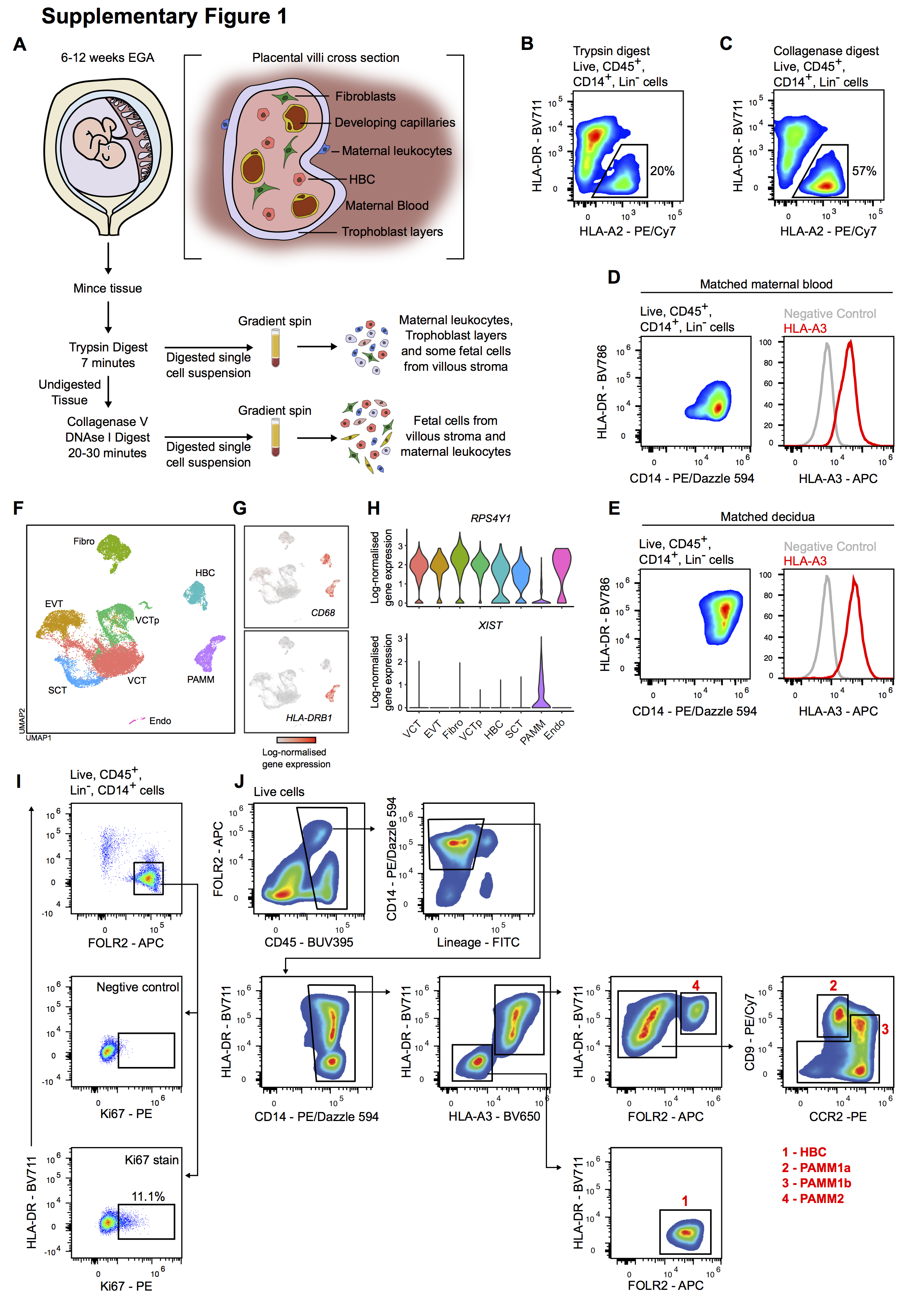

### Figure S2

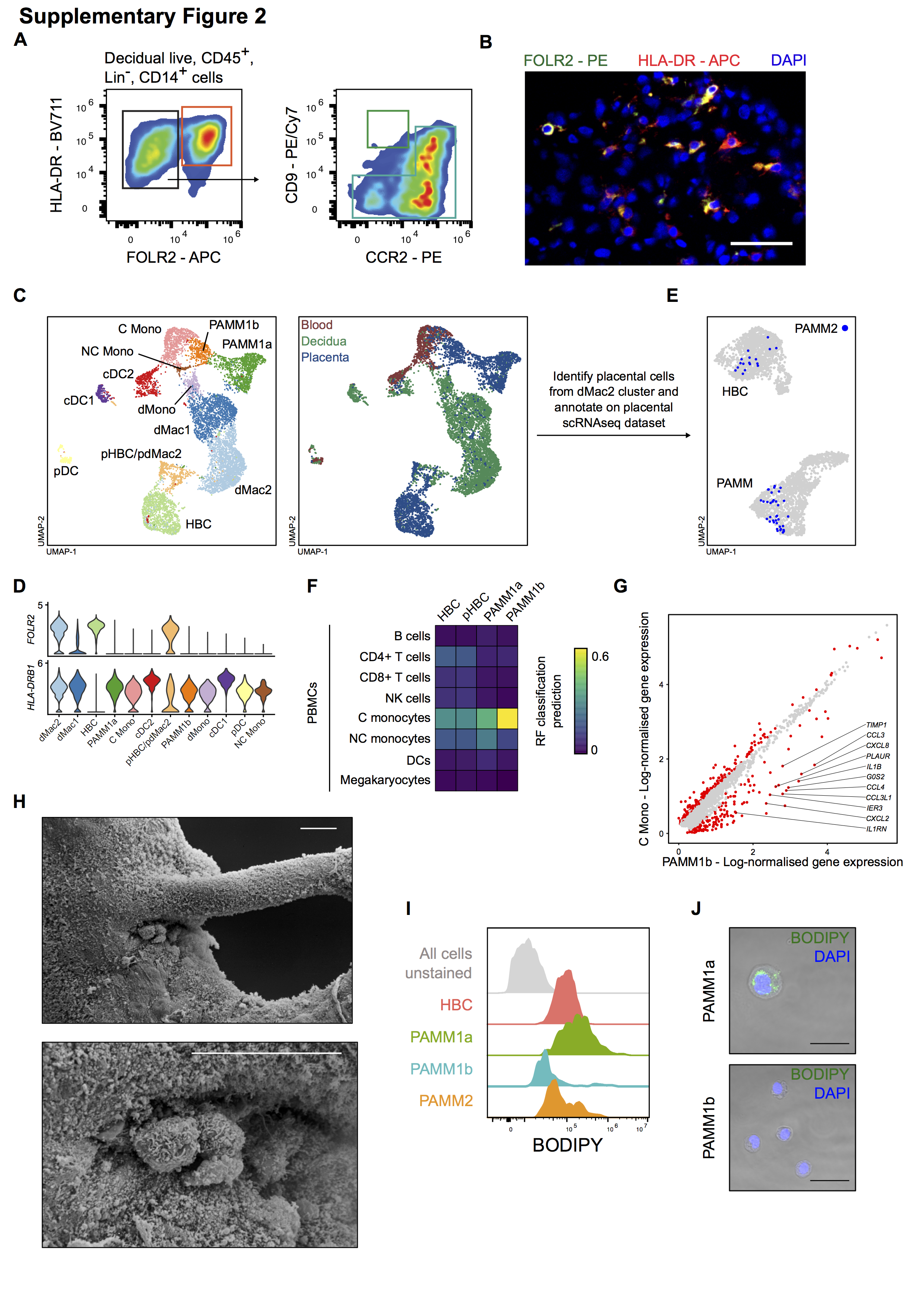

### Figure S3

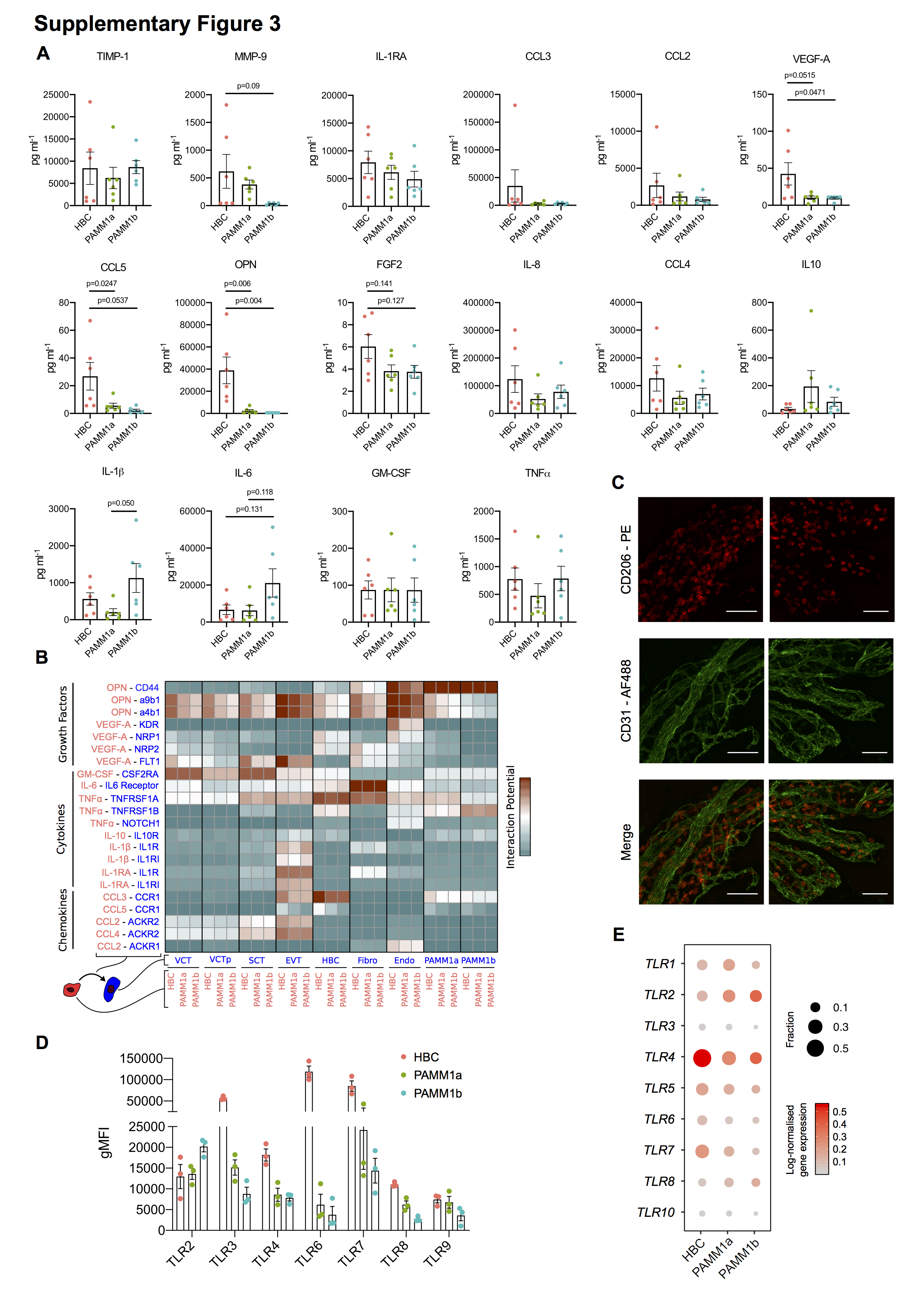

### Figure S4

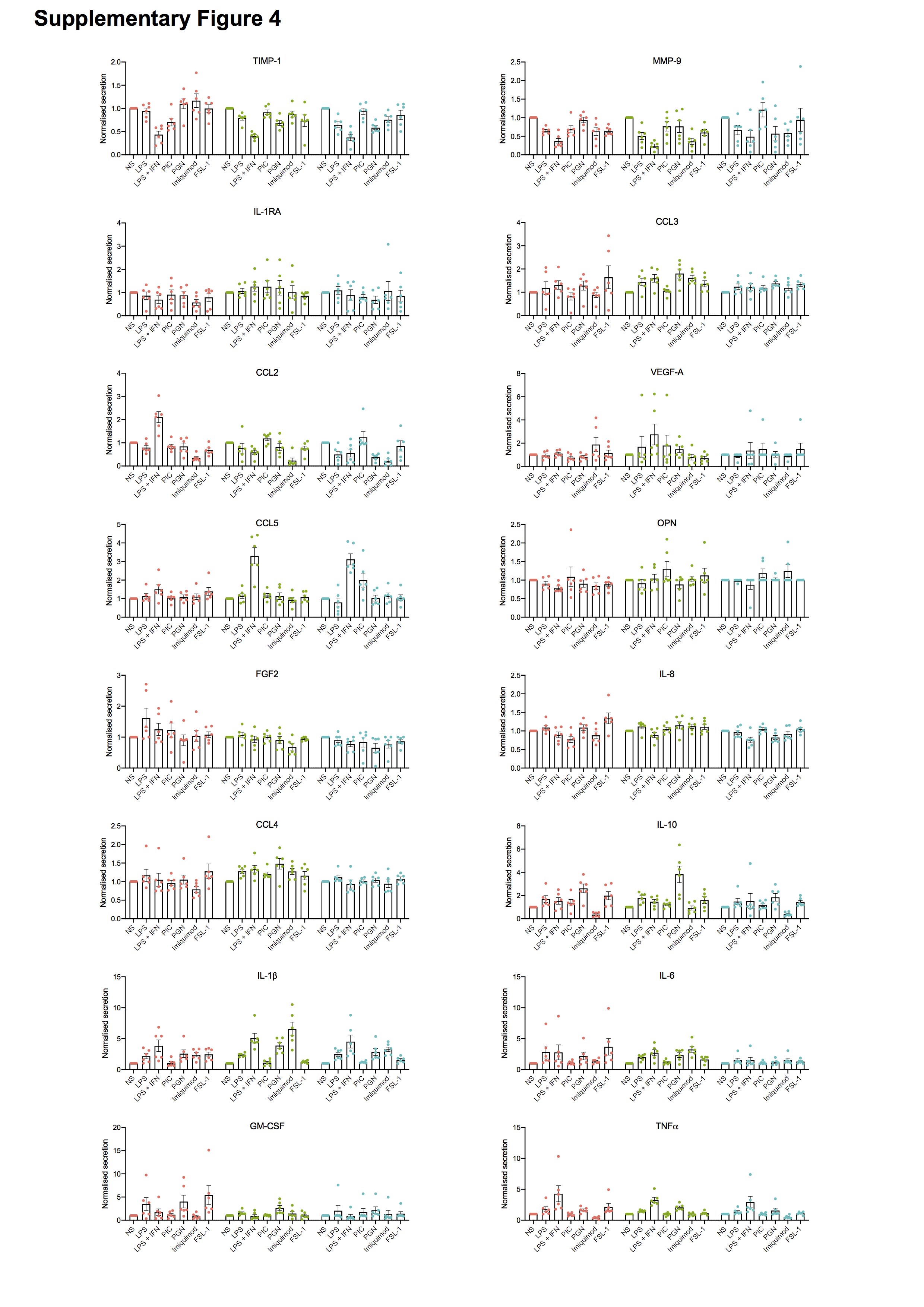

### Figure S5

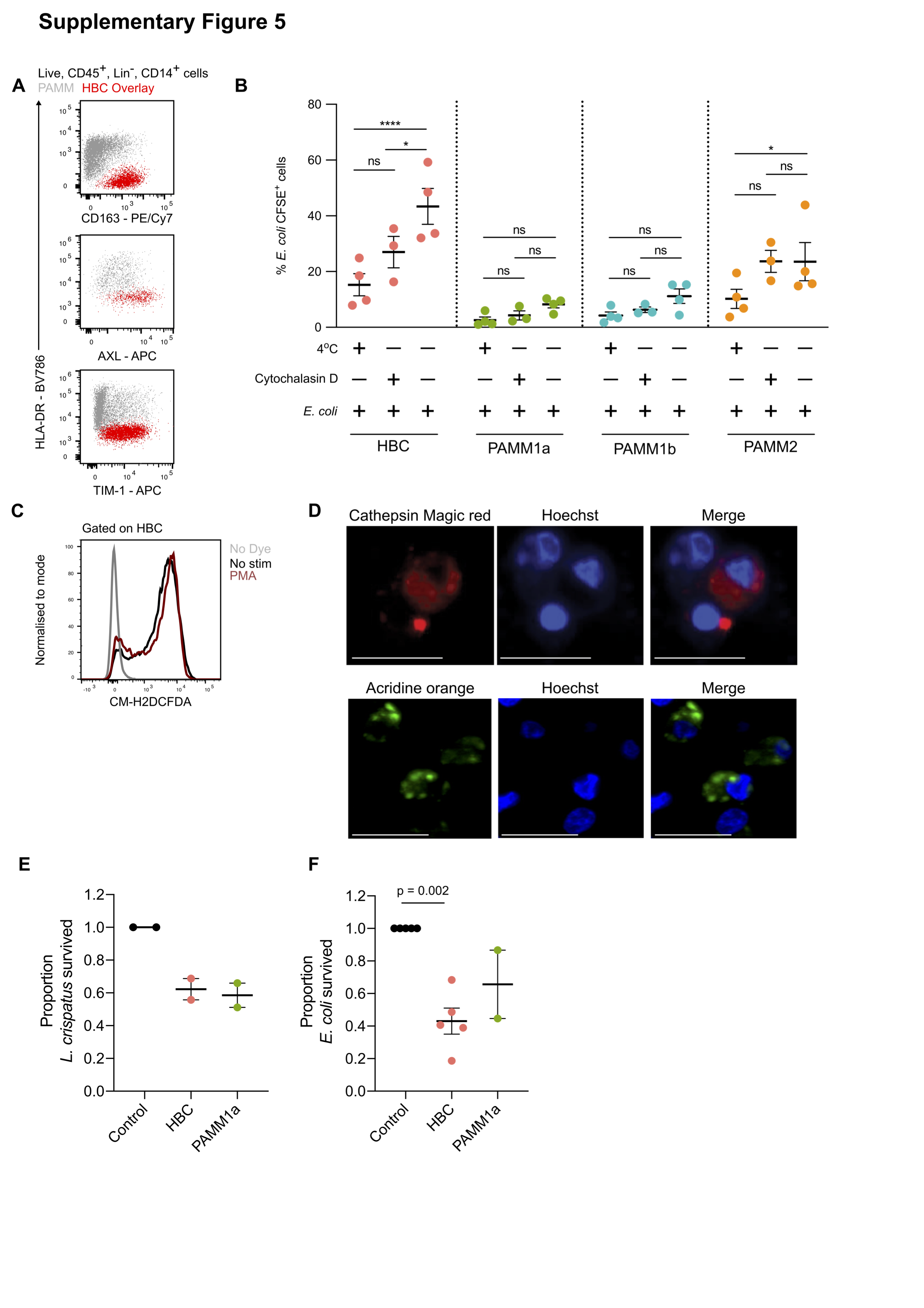
